# Ecological determinants of disease and immunity in myelodysplastic syndromes

**DOI:** 10.64898/2026.05.05.720208

**Authors:** Robert F. Stanley, Beatrice D. Zhang, Kimon V. Argyropoulos, Pu Zhang, Maxim Maron, Brianna Gipson, Chloe Park, Kenyon Weis, Alexander M. Lewis, Zoe Katsamakis, Charlotte Wishnack, M. Adriana Cuibus, Ning Fan, Karen Zhao, Kenton Wu, Catherine Snopkowski, Joshua Weinreb, Jeetayu Biswas, Matthew Zatzman, Nathan Aleynick, Leonardo Boiocchi, Megan So-Young Lim, Roni Tamari, Jonathan U. Peled, Gunjan Shah, Andrew Moorman, Yuval Elhanati, Eric Rosiek, Mikhail Roshal, Ahmet Dogan, Umesh Bhanot, Eytan M. Stein, Sarah Samorodnitsky, Ronan Chaligne, Marcel R.M. van den Brink, Stephen Martis, Benjamin D. Greenbaum, Omar Abdel-Wahab, Susan DeWolf

## Abstract

Myelodysplastic syndromes (MDS) are clonal hematopoietic malignancies characterized by ineffective hematopoiesis, dysplastic morphology, and risk of progression to acute myeloid leukemia. While genomic alterations intrinsic to malignant MDS disease-initiating cells have been well-characterized, molecular assessment of the bone marrow *in situ* has been limited. Here we present single cell spatial assessment of gene expression, T cell receptors, as well as MDS-defining mutations and RNA isoforms within fixed, decalcified human bone marrow core biopsies (41 MDS, 15 controls) paired with single cell immunogenomic analysis of bone marrow aspirates (35 MDS, 6 controls). Bone marrow spatial analyses of >7.47×10^6^ cells identified hematopoietic and non-hematopoietic populations not readily captured in dissociated tissue. We developed computational methods to compare ecological niche structures, revealing enriched hematopoietic niches and reorganization of T cell immunity in MDS patient bone marrow. *In situ* genotyping of mutant cells revealed increased TGFβ expression in malignant megakaryocytes suppressing local T cell cytotoxicity. By contrast, TGFβ signaling was disrupted in mutant cells due to aberrant splicing of multiple TGFβ signaling components. This study provides a spatially resolved landscape of human MDS bone marrow and uncovers mechanisms by which malignant cells simultaneously promote intrinsic clonal persistence while rewiring the microenvironment for immune escape.

## INTRODUCTION

Myelodysplastic syndromes (MDS) are a heterogenous group of clonal myeloid blood cancers arising from hematopoietic stem cells within the bone marrow. MDS is characterized by morphologic dysplasia of hematopoietic cells, ineffective hematopoiesis, and risk of transformation to acute myeloid leukemia^1^. Genomic studies of human MDS have largely focused on hematopoietic cell-intrinsic analyses of disaggregated cells from marrow aspirates. These studies have been instrumental in defining the genomic landscape of MDS and have revealed that among cancers, MDS is uniquely characterized by a high frequency of mutations in epigenetic regulatory proteins and RNA splicing factors^2,3^. Clinical evaluation via bone marrow core and aspirate biopsy is essential to adequately quantitate and characterize dysplasia and identify cytogenetic abnormalities and somatic mutations that are central to MDS diagnosis, classification, and prognostication^4,5^. Beyond hematopoietic cell-intrinsic genomic alterations, several lines of evidence indicate that the bone marrow microenvironment is a key contributor to MDS pathogenesis and related myeloid clonal disorders^6,7^. For example, foundational studies in animal models have revealed that alterations in the stroma of MDS can functionally drive MDS development^8,9^. Furthermore, compelling studies in both patients and preclinical models have identified that the MDS bone marrow is associated with increased inflammation and aberrant immunity, primarily in the innate compartment^10–13^, whereas data regarding T cell immunity are limited^14^. The link between MDS and aberrant immune activation is underscored by the well-established connection between MDS and a variety of autoimmune conditions^15^.

Despite the importance of the bone marrow microenvironment to MDS pathogenesis, high resolution studies of human bone marrow in an *in situ* unperturbed state have been limited. To date, spatial studies of human bone marrow in MDS have utilized protein-based detection methods unable to characterize molecular cell phenotypes at single cell resolution^16–18^. Furthermore, the application of spatial transcriptomics to bone marrow tissue has been hampered by technical challenges of performing molecular studies on acid-decalcified marrow cores. As such, the spatial architecture of the immune microenvironment in human MDS has not been studied at single cell resolution. Utilizing spatial tools to improve our understanding of cell-cell interactions in the MDS marrow including the relationships among stromal populations, immune subsets, and malignant clones is fundamental to dissecting the role of the marrow microenvironment in disease pathogenesis, including understanding features of aberrant immunity in MDS patients.

Here, we characterize the human MDS bone marrow microenvironment and adaptive immune landscape utilizing a multiomic approach combining spatial and single cell molecular analyses. We applied EDTA decalcification to enhance preservation of nucleic acids for single cell spatial transcriptomic profiling of human bone marrow core biopsies with a marrow and T cell focused >400-plex gene expression panel (Xenium, 10x Genomics). Further, we developed a pair of computational tools for the ecological analysis of spatial transcriptomics data. The first we term “SAND” (**S**patial **A**djacency **N**etwork **D**etection), which identifies discrete, biologically-relevant structures within tissues like bone marrow that lack typical spatial landmarks. The second, “DUNE” (**D**iscovery by **U**nsupervised **N**iche **E**nrichment), defines enriched local niches and enables unsupervised comparison of niche enrichment between samples. In parallel, we performed single cell protein, RNA, and TCR sequencing of MDS patient bone marrow aspirates, capturing the diversity of adaptive immune phenotypes and T cell clonality across MDS. TCR sequencing of bone marrow aspirates enabled single cell spatial identification of clonally expanded CD8^+^ T cells in MDS bone marrow *in situ* using custom CDR3 targeting probes. Moreover, we designed bespoke probes to simultaneously capture MDS-defining somatic mutations in the core RNA splicing factors SF3B1 and U2AF1 along with aberrant mRNA isoforms expressed selectively in SF3B1 mutant cells. These efforts uncovered mechanisms of clonal selection for MDS cells driven by both mutant cell-autonomous and non-cell-autonomous effects within the local microenvironment.

## RESULTS

### Comprehensive single cell spatial and multiomic interrogation of human MDS

To study the MDS bone marrow microenvironment at single cell resolution *in situ*, we performed spatial transcriptomic analysis on formalin fixed, EDTA-decalcified human bone marrow core biopsies (see **Methods**), enabling interrogation of hundreds of genes simultaneously in the context of their cellular localization (**Fig. 1a**). We built a >400 gene panel for the Xenium platform (10x Genomics) combining probes for 377 genes from the Xenium Human Multi-Tissue and Cancer panel with 56-100 custom probes to define critical bone marrow populations, including detailed lymphocyte subsets (**Supplementary Table 1**). We applied this panel to core biopsies from 41 patients with newly diagnosed untreated MDS, 15 aged-matched healthy controls, and 5 MDS patients in remission after hematopoietic cell transplantation (HCT), capturing >5.7 million high-quality single cells (clinical details in **Supplementary Table 2, Fig. 1b**). We prioritized early-stage newly diagnosed, untreated MDS in order to capture initial remodeling of the bone marrow microenvironment prior to disease progression. The median age of MDS patients was 70 years old, and the majority were diagnosed with MDS with low-blasts (MDS-LB, n=24) including a subset of MDS with low blasts and SF3B1 mutation (MDS-*SF3B1*, n=7); we also studied 12 patients with MDS with increased blasts 1 (IB1) and five patients with MDS with increased blasts 2 (IB2). Our control cohort was comprised of 10 lymphoma-negative staging core bone marrow biopsies and five healthy bone marrow transplant donors. Finally, we also analyzed bone marrow biopsies from five patients with MDS in remission for over 2 years after allogeneic-HCT with no clinical evidence of disease at time of biopsy.

**Fig. 1:**
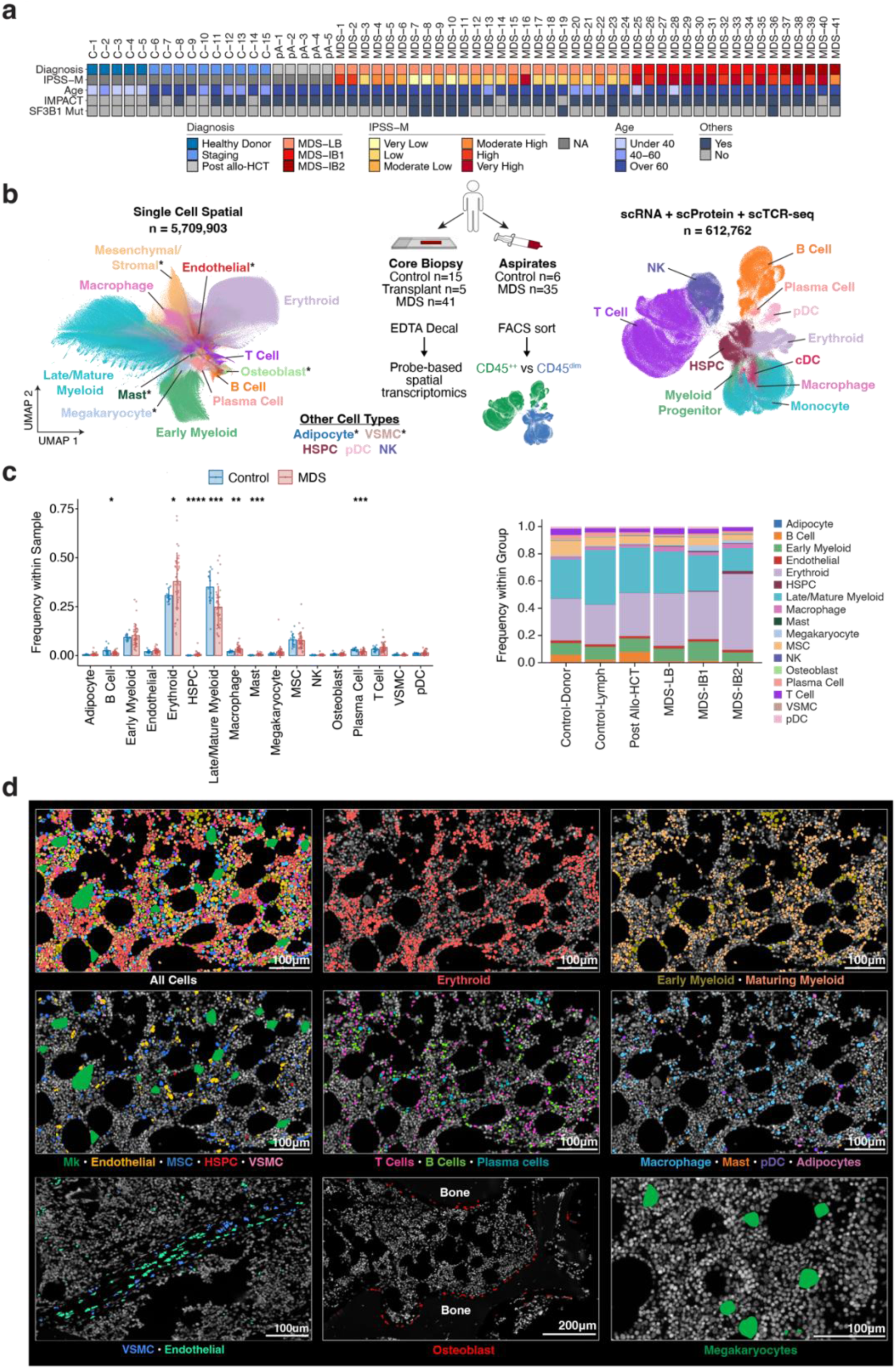
Comprehensive single cell spatial and multiomic analysis of human MDS bone marrow. **a,** Oncoprint of patient bone marrow samples used for single cell spatial analysis. From top to bottom: Patients were diagnosed according to WHO 2022 classification^72^. Molecular International Prognostic Scoring System (IPSS-M) was used to determine MDS risk category^4^. The age of each patient at sample collection is indicated. Samples subject to Integrated Mutation Profiling of Actionable Cancer Targets (IMPACT) testing^53^, and samples with SF3B1 mutations are annotated. **b,** Schematic of overall study design. Healthy control (n=15), post allogenic stem cell transplantation (n=5), and newly diagnosed and/or untreated MDS (n=41) patient bone marrow core biopsies were EDTA decalcified, and single cell spatial transcriptomics was performed. Bone marrow aspirates from healthy controls (n=6) and MDS patients (n=35) were flow sorted into CD45^++^ and CD45^dim^ populations, hashtagged, and CITE-seq and TCR-seq was performed. UMAP representations of 5,709,903 single cells from spatial analysis of bone marrow patient samples (left) and 612,762 single cell transcriptomes of bone marrow aspirates (CITE-seq, right) are colored by cell type. Asterisks indicate cell populations not captured by single cell bone marrow aspiration-based techniques. **c,** Bar plots (left) comparing cell type frequencies between control (donor and staging marrows) and MDS single-cell spatial samples. P-values were calculated by Wilcoxon rank-sum test: *p ≤ 0.05, **p ≤ 0.01, ***p ≤ 0.001, ****p ≤ 0.0001. Stacked bar plots (right) of cell type frequencies in spatial samples stratified by sample type. **d,** Representative images of bone marrow cell populations highlighting 16 different cell types. Top and middle rows: Representative images of different cell populations highlighted by color. Bottom row: Representative images of key cell populations not easily captured by aspiration-based techniques. TCR, T cell receptor; HSPC, hematopoietic stem and progenitor cell; NK, natural killer cell; VSMC, vascular smooth muscle cell; pDC, plasmacytoid dendritic cell; cDC, conventional dendritic cell; MSC, mesenchymal stromal cell; Mk, megakaryocyte.

In parallel to our spatial analyses, we performed single cell gene expression, surface marker (CITE), and T cell receptor (TCR) sequencing on 58 bone marrow aspirates (single cell suspensions) from 35 untreated MDS patients (clinical details **Supplementary Table 3**), nine of which had longitudinal samples, and six control subjects (>6×10^5^ cells and >1.55×10^5^ TCR, **Extended Data Fig. 1a**). Of note, 10 aspirates were acquired at the sample time of a core biopsy (**Supplementary Table 3**), enabling direct comparison to spatial samples.

Our workflow for preprocessing spatial data included cell segmentation, quality control, and subsequent cell typing (details in **Methods**, **Extended Data Fig. 1b**). We manually annotated all cell populations using established lineage markers, delineating >20 distinct bone marrow cell populations including T and B cell immune subsets (**Extended Data Fig. 1c-e**). Importantly, spatial profiling of core marrow biopsies allowed for detection of cell types with established or putative relevance in MDS pathology but not readily captured by aspiration-based bone marrow analyses including megakaryocytes, endothelial cells, mesenchymal stromal cells (MSC), vascular smooth muscle cells, adipocytes, mast cells, and osteoblasts (**Fig. 1b-d** and **Extended Data Fig. 2a-b**)^8,9,19–21^.

Whereas most solid organs are organized into compartments such as epithelial layers, lumens, cortices, and other clearly demarcated microanatomical zones, the lack of morphologically organized structures in human bone marrow compared to solid tissues presented an analytic challenge for defining the physical distribution of cell types. To this end, we developed a highly generalizable graph-based clustering method (**SAND**: **S**patial **A**djacency **N**etwork **D**iscovery) to spatially define local networks consisting of predefined features (such as clusters of lymphocytes), enabling the identification of biologically relevant marrow structures (see **Methods**). In brief, SAND constructs a spatial neighborhood graph which is pruned to include only nodes representing feature categories of interest (for example, T cells and B cells) and subsequently identifies connected components as clusters. We applied this approach to identify clusters of over-segmented megakaryocytes and refine their segmentation. Because of their multinucleated structure and dysplastic morphology in MDS, naïve cell segmentation calls frequently divided a single megakaryocyte into >10 different individual cells (**Extended Data Fig. 2c-d)**.

With respect to overall bone marrow composition, frequencies of many cell types were similar across all samples (**Extended Data Fig. 2e**); however, MDS samples had higher frequencies of erythroid cells and hematopoietic stem and progenitor cells (HSPCs) (**Fig. 1c**), an effect mostly attributable to samples from patients with advanced MDS (IB1/IB2) (**Extended Data Fig. 3a**). Consistent with prior studies^25^, we found a paucity of B cells compared to controls (-0.99 log_2_ fold-change, p=0.012) (**Fig. 1c**), making T cells the most abundant lymphocyte subtype in MDS marrow (**Extended Data Fig. 3a-b**).

### Unsupervised analyses reveal hematopoietic niche structures enriched in MDS

Our expansive cohort of control and MDS samples of varying stages enabled sample-level comparison of bone marrow structure. To do so, we developed a complementary computational method to quantify the enrichment of local ecological niches by sample (DUNE: **D**iscovery by **U**nsupervised **N**iche **E**nrichment). Each niche is defined as the cell type composition of a cell and its k-nearest neighbors (**Fig 2a**). The enrichment score for a niche in a sample is computed by normalizing the observed count of the niche by the expected count computed from global sample cell type frequencies (see **Methods**). Principal component analysis (PCA) of filtered niche enrichment scores revealed that bone marrow biopsies from adults without evidence of MDS (both healthy donor subjects and bone marrows acquired for lymphoma staging, as well as post-transplant samples from patients in remission from MDS) clustered closely together (see **Methods**, **Fig. 2b**). Samples from individuals without MDS also formed a cluster in PCA space based on cell type proportions (**Extended Data Fig. 3c**). These data indicate that the spatial architecture of the bone marrow in patients in long-term remission from MDS post-allogeneic transplantation resembles that of healthy individuals. In contrast, samples from patients with MDS were relatively scattered in niche enrichment PCA space, indicating spatial heterogeneity among MDS samples (**Fig. 2b**). Unsupervised clustering of samples on filtered niche enrichment scores revealed a cluster exclusively composed of MDS samples including all MDS-IB2 samples and 46% of all MDS samples (**Fig. 2c, Extended Data Fig. 3d**).

**Fig. 2:**
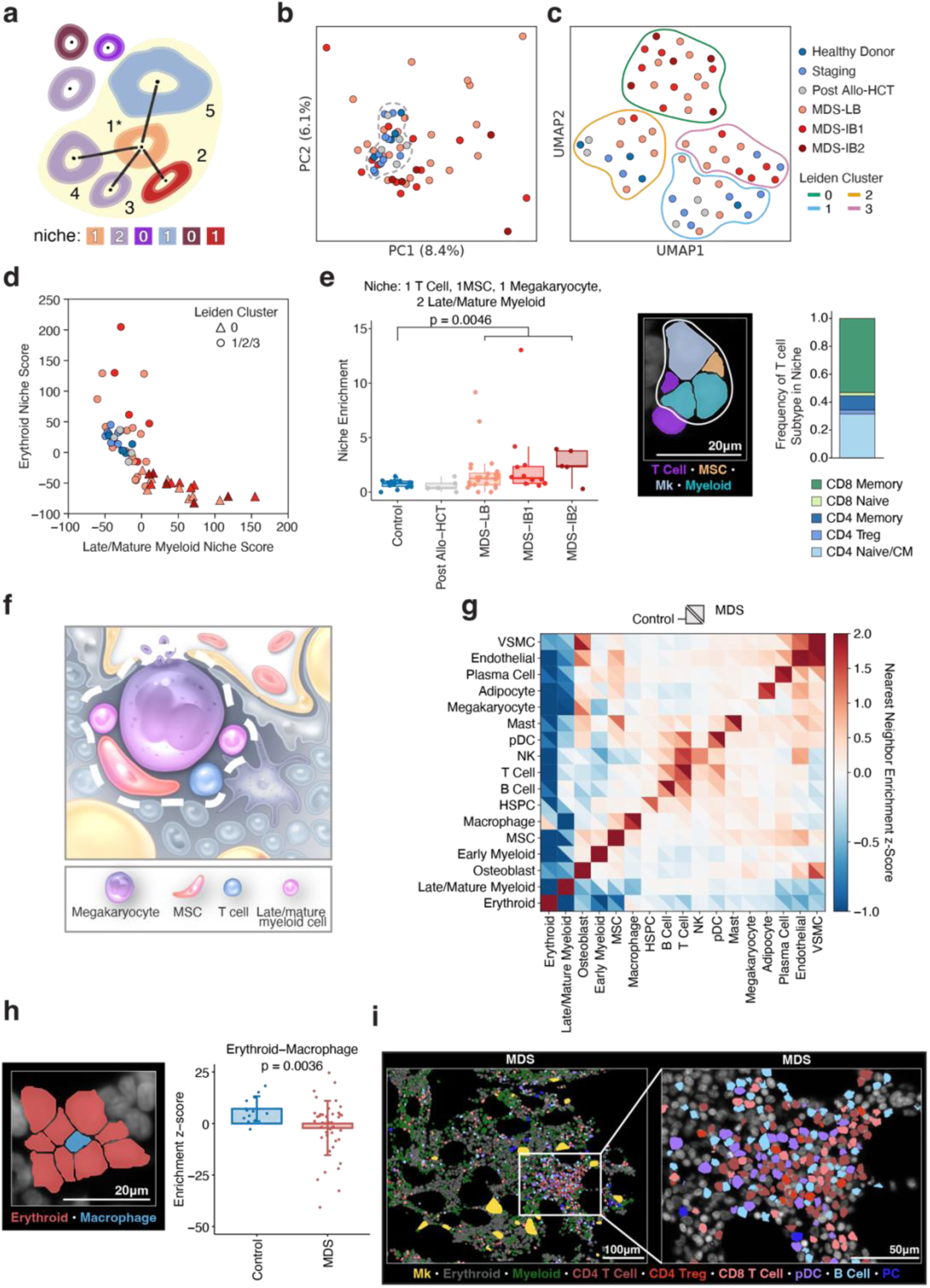
Altered spatial distribution and cell-cell relationships in MDS bone marrow. **a,** Schematic of a cell niche composed of a central cell and its four nearest neighbors (black lines). Black dots represent the centroid of each cell. The niche is identified by a vector of cell type counts. **b**, Principal component analysis visualization of samples computed on niche enrichment scores colored by diagnosis. The dotted line encircles all non-MDS samples. **c,** UMAP visualization of samples computed on niche enrichment scores. Samples are colored by diagnosis and colored lines partition Leiden clusters. **d,** Summary plot of differential niche enrichment analysis. Differentially enriched niches of Leiden cluster 0 were first determined. Then, the Late/Mature Myeloid niche score for each sample was computed by taking the sum of niche enrichment scores across differentially enriched niches containing ≥1 Late/Mature Myeloid cell. The Erythroid niche score is similarly computed as the sum of enrichment scores of differentially enriched niches containing ≥1 Erythroid cell. Each point is colored by diagnosis, and samples assigned to Leiden cluster 0 are denoted by triangles. **e**, Boxplot (left) of niche enrichment scores of a differentially enriched niche consisting of one T cell, one mesenchymal stromal cell (MSC), one megakaryoctye (Mk), and two Late/Mature Myeloid cells. The p-value comparing the enrichment scores of this niche between control and MDS samples is calculated with a Wilcoxon rank-sum test. A representative image of this niche is displayed (center), and subtype composition of T cells occurring in this niche is displayed as a stacked bar graph (right). **f**, Schematic of an MDS-enriched cellular niche composed of one T cell, one MSC, one megakaryocyte, and two late/mature myeloid cells. This niche is a feature of the late/mature myeloid axis and increases with disease progression. **g**, Neighborhood enrichment z-score heatmap between pairs of bone marrow cell populations (squares) from control (bottom triangle) and MDS (top triangle) patients. Red indicates positive enrichment (positively associated, > 0), blue indicates negative enrichment (negatively associated, <0). **h**, Representative image of an erythroid island (left). Bar plot (right) showing the average neighborhood enrichment z-score between erythroid and macrophage populations in control and MDS samples. Error bars represent standard deviation. P-value was calculated by Wilcoxon rank-sum test. **i**, Left: Representative image of MDS bone marrow with white box highlighting an MDS immune aggregate. Right: Higher magnification of MDS immune aggregate. MDS with low blasts (MDS-LB); MDS with increased blasts 1 (MDS-IB1); MDS with increased blasts 2 (MDS-IB2). pDC, plasmacytoid dendritic cell; Mk, megakaryocyte; Treg, regulatory T cell; PC, plasma cell; mesenchymal stromal cell, MSC.

To dissect which ecological niches contribute to this clustering, we performed differential niche enrichment analysis between samples residing in the MDS-exclusive cluster and all other samples. In doing so, we discovered two primary axes of variation that drive differentiation of MDS samples from each other and from controls (**Fig. 2d**, **Extended Data Fig. 3e**). The first is along an erythroid-enriched niche axis (y axis, **Fig. 2d**), which defines controls and MDS samples that do not reside in the MDS-exclusive cluster. The second is a late/mature myeloid-enriched axis (x axis, **Fig. 2d**) which defines the MDS exclusive cluster that includes all IB2 samples. An example of such a niche is one composed of one T cell, one MSC, one megakaryocyte, and two late/mature myeloid cells. The enrichment score of this niche is not only significantly higher in MDS samples compared to controls but also increases with disease progression (**Fig. 2e-f**). Further inspection of T cell subtypes revealed that the majority of T cells present in this niche are CD8 memory cells with upregulation of *CCL5, GZMA,* and *GZMK* (**Fig. 2e and Extended Data Fig. 3f**).

To interrogate pairwise cell-cell relationships driving differences between human MDS marrow and controls, we performed a nearest neighbor analysis for each major cell type.

Specifically, we identified the five nearest neighbors for each cell population and derived a neighborhood enrichment z-score^22^ for all pairs of cell types which were then visualized in a heatmap stratified by MDS and control marrows (**Fig. 2g, Extended Data Fig. 3g**). Erythroid cells were positively associated with other erythroid cells, validating the identification of erythroid islands characteristically seen by traditional H&E staining, and negatively associated with other cell types (**Fig. 2g**) with the degree of negative association typically greater in MDS when compared to control samples. One interesting exception is that there was a positive association between macrophages and erythroid cells in controls while the association was negative in MDS. This has been described previously^23,24^ and was supported by our previous niche enrichment analysis (**Fig. 2h, Extended Data Fig. 3h**). Mature myeloid cells exhibited a similar pattern of self-association, with the degree of negative association with other cell types being greater in controls than in MDS samples. Collectively, this pairwise analysis suggests that subtle coordinated shifts in cell-cell associations drive the differences in MDS and controls described in **Figs. 2b-d**. ^23,24^

### Remodeling of adaptive immunity in human MDS bone marrow

Focusing on the relationship between adaptive immune subsets, we performed pairwise cell type neighborhood analysis of all lymphocyte subsets. This analysis identified distinct differences in the distribution of T and B cell populations in MDS compared to controls (**Extended Data Fig. 3i**). Our nearest neighbor analysis also suggested physical proximity of clusters of immune cells comprised of T-, B-, natural killer (NK), and plasmacytoid dendritic cells (pDCs) within the MDS marrow (**Fig. 2g**). We confirmed the presence of these structures via spatial visualization in individual bone marrow cores. This revealed “immune aggregates,” spatially localized lymphocyte populations sometimes reminiscent of tertiary lymphoid structures (TLS), distributed throughout the marrow (**Fig. 2i**).

We applied SAND to identify individual immune aggregates throughout the marrow of all patients and elucidate their cellular architecture. We classified any spatial cluster of lymphocytes with ≥10 T cells and ≥20% T cell composition (see **Methods**, **Extended Data Fig. 4a**) as an aggregate. While there was no significant difference in the size of immune aggregates between MDS and controls (**Extended Data Fig. 4b-c)**, lymphocytes in MDS marrows were enriched within immune aggregate regions (**Fig. 3a-b**). This was also reflected by the increased association between lymphocytes in MDS compared to controls from our previous nearest neighbor analysis (**Fig. 2g**). Furthermore, immune aggregates in MDS had distinct compositions and organization compared to controls. Lymphoid aggregates in control marrows were composed of a mixture of CD4 T, CD8 T, and B cells. In contrast, small immune aggregates in MDS (<100 cells) were enriched in CD4 and CD8 T cell populations but depleted in B cells and larger MDS aggregates (≥100 cells) were enriched in CD4 T and B cells but depleted in CD8 T cells (**Fig. 3c** and **Extended Data Fig. 4d-e**), with further distinctions across T and B cell subsets (**Extended Data Fig. 4f).**

**Fig. 3:**
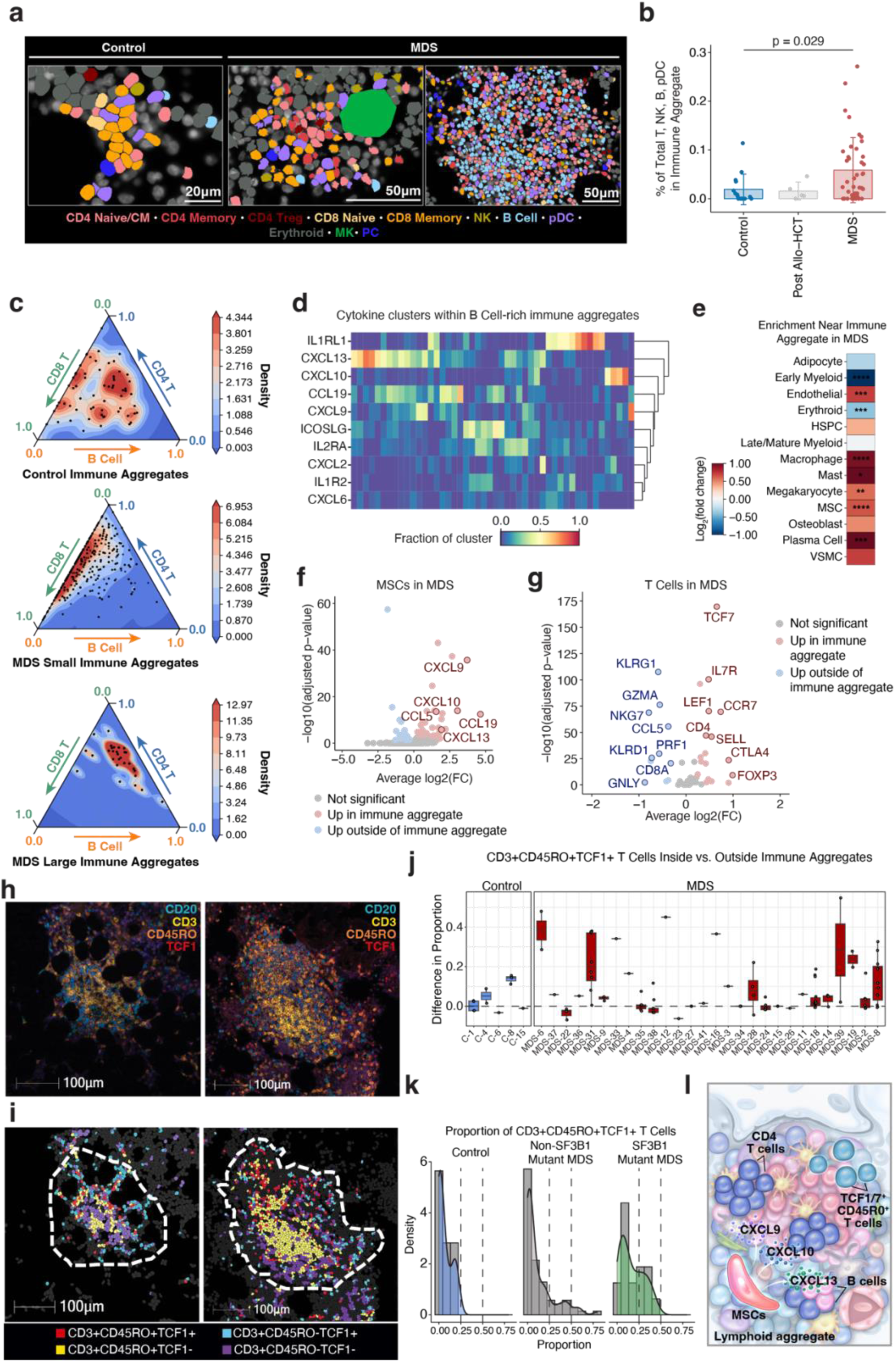
Distinct lymphocyte and cytokine composition of MDS immune aggregates with enrichment of *TCF7*/TCF1^+^ stem-like memory cells. **a,** Representative images of control and MDS bone marrow immune aggregates. Left: Small control immune aggregate (<100 cells). Middle: Small MDS immune aggregate (<100 cells). Right: Large MDS immune aggregate (≥100 cells). **b**, Bar plot of proportion of T, NK, B, and pDC cells located within immune aggregates in each sample. Each bar represents the average, and error bars represent standard deviation. P-value was calculated by Wilcoxon rank-sum test. **c**, Ternary plots showing the frequency of B, CD4 T, and CD8 T cell populations in control immune aggregates (top), small (<100 cells) MDS immune aggregates (middle), and large (≥100 cells) MDS immune aggregates (bottom). **d**, Heatmap displaying the composition of cytokine clusters (columns) colocalized with B Cell-rich (≥33% B Cell) immune aggregates. **e**, Heatmap displaying spatial enrichment near immune aggregates for each cell type (excluding T, B, NK, and pDCs). Enrichment is calculated by computing the average log2(fold change) between the frequency of each cell type out of all cells that are near (≤15µm from a T, B, NK, or pDC within an immune aggregate) versus far from immune aggregates over all MDS samples containing immune aggregates. Asterisks denote p-values computed from paired Wilcoxon signed-rank test: *p ≤ 0.05, **p ≤ 0.01, ***p ≤ 0.001, ****p ≤ 0.0001. **f**, Volcano plot of differential gene expression between mesenchymal stromal cells (MSCs) near immune aggregates (≤15 µm from a T, B, NK, or pDC within an immune aggregate) versus MSCs far from immune aggregates in MDS samples. **g**, Volcano plot of differential gene expression between T cells within immune aggregates vs T cells outside of immune aggregates in MDS samples. **h**, Representative multiplex immunofluorescence photomicrographs of immune aggregates; CD20 (cyan), CD3 (yellow), CD45RO (orange), TCF1 (red). **i,** Representative masks of different T cell immunophenotypes in aggregates from **h;** CD3^+^CD45RO^+^TCF1^+^ (red), CD3^+^CD45RO^+^TCF1^-^ (yellow), CD3^+^CD45RO^-^TCF1^+^ (cyan), CD3^+^CD45RO^-^TCF1^-^ (purple). **j,** Difference in the proportion of CD3^+^CD45RO^+^TCF1^+^ T cells in each immune aggregate vs. scatter regions within each core marrow biopsy (MDS marrows; p=0.001; healthy control marrows; p=0.607). **k**, Distribution of the proportion of CD3^+^CD45RO^+^TCF1^+^ subsets out of total T cells across all immune aggregates shows a statistically significant difference when comparing MDS with SF3B1 mutations versus SF3B1 wild-type marrows (Kruskal-Wallis rank sum test, p=0.016). Vertical lines denote frequencies of CD3^+^CD45RO^+^TCF1^+^ subsets at 25% and 50%, respectively. **l**, Schematic of an MDS lymphoid aggregate showing MSCs, CD4 T cells and TCF1/7^+^CD45R0^+^ T cells and B cells. Local production of CXCL9, CXCL10, and CXCL13 by MSCs supports coordinated recruitment and retention of lymphoid populations to establish a structured localized immune microenvironment. NK, natural killer cells; pDC, plasmacytoid dendritic cell; Mk, megakaryocyte; Treg, regulatory T cell; PC, plasma cell; mesenchymal stromal cell, MSC; CD4 naïve/CM, CD4 naïve/central memory.

Principal component analysis of pairwise cell type neighborhood enrichment scores within immune aggregates and neighboring cells revealed that larger MDS immune aggregates had increased immune cell density and structure compared to smaller MDS aggregates and aggregates in controls (**Fig. 3c** and **Extended Data Fig. 4g-h**). Given that specific chemokines and cytokines have been reported to drive the formation of immune aggregates in other cancer types (*CXCL9*, *CXCL10*, *CXCL13*, and *CCL19* among others^25–27^), we applied SAND to define cytokine and chemokine clusters across spatial bone marrow data (**Extended Data Fig. 5a**). This revealed that larger cytokine clusters colocalized with immune aggregates (**Extended Data Fig. 5b)** and an association between chemokine and lymphocyte compositions of aggregates (**Fig. 3d**, **Extended Data Fig. 5c**). Furthermore, transwell assays with primary MDS bone marrow mononuclear cells demonstrated the capacity of these chemokines to drive the migration of specific lymphocyte subsets (**Extended Data Fig. 5d**). Analysis of the cells surrounding immune aggregates in MDS revealed enrichment of myeloid cells and stromal cells including MSCs, which were not increased around control aggregates (**Fig 3e, Extended Data Fig. 5e**). Moreover, differential gene expression of MSCs near versus far from MDS immune aggregates revealed increased expression of TLS-associated chemokines (**Fig. 3f**).

In MDS but not controls, we found distinct changes in composition of lymphocyte subsets within immune aggregates compared to areas outside immune aggregates: in MDS, we found an increase in CD4 naïve and central memory T cells as well as regulatory T cells (Tregs), whereas CD8 memory T cells were relatively increased outside of these aggregates (**Extended Data Fig. 5f**). Analysis of differentially expressed genes in memory T cells within and outside MDS immune aggregates revealed increased expression of *TCF7*, which encodes the transcription factor TCF1 and is linked to T cell stemness^28^ (**Fig. 3g** and **Extended Data Fig. 5g**) and has been associated with immune aggregate and TLS formation in other cancers^29,30^. To validate the composition and phenotype of lymphocytes within immune aggregates at the protein level, we developed a multiplexed immunofluorescence (IF) panel comprised of CD20 and CD3 to define B and T cells, respectively, CD45RO to identify memory T cells, and TCF1. We applied this panel to 43 bone marrow core biopsies (7 controls, 36 MDS) that also underwent spatial transcriptomic evaluation. Analysis was restricted to samples containing at least one immune aggregate with >50 T and/or B cells, yielding 33 evaluable biopsies (5 controls, 28 MDS; see **Methods**), which were present in both MDS and control marrows (**Fig. 3h-i** and **Extended Data Fig 6a-c**). TCF1 expression within immune aggregates was not significantly different between controls and MDS patients; however, patients with MDS-LB tended to have greater *TCF7*/TCF1 expression by transcriptional and protein analysis compared to patients with MDS-IB1/2 (**Extended Data Fig. 6d-e**). When TCF1 expression was analyzed across T cell subsets, T cells within immune aggregates included a higher frequency of CD45RO+TCF1+ stem-like memory T cells compared to regions outside aggregates, a pattern not observed in controls (**Fig. 3j** and **Extended Data Fig. 6f-g**). Notably, a subset of MDS immune aggregates across disease states contained >25% stem-like memory T cells, a feature absent in controls and more pronounced in SF3B1-mutant MDS (**Fig. 3k** and **Extended Data Fig. 6h-i).** Taken together, these data demonstrate the utility of SAND in resolving spatial immune architecture in the bone marrow and define the lymphocyte and cytokine composition of MDS immune aggregates, including enrichment of *TCF7*/TCF1^+^ stem-like memory T cells (**Fig. 3l**).

### Tracking CD8 T cells across space and time in MDS

Given the altered spatial distribution of lymphocytes identified in MDS, we performed whole transcriptome single cell analysis of bone marrow aspirates to more comprehensively interrogate cell composition in MDS, including investigation of T cell repertoire diversity. We compared the cell composition and differential abundance of immune subsets across untreated MDS patients (n=34), MDS patients treated with azacitidine (n=3; all with persistent disease at time of evaluation), post-allogeneic transplant samples (n=10), and healthy control morrow aspirates (n=6) (**Fig. 4a**, **Extended Data Fig. 7-8**). Consistent with prior studies^31^ and our spatial data, immature and mature B cell populations were significantly lower in MDS compared to controls (p=0.0006, p=0.021 respectively), resulting in a relative enrichment in T and NK cells in MDS (**Extended Data Fig. 9a**). Notably, there was no significant difference in the frequency of CD4 T cell subsets in MDS compared to controls, whereas among CD8 subsets, *GZMB^+^*(granzyme B) T cells were significantly enriched in MDS (p=0.0045) (**Extended Data Fig. 9b**). These *GZMB^+^* CD8 memory T cells expressed high levels of perforin mRNA as well as protein-level expression of CD45RA, TIGIT, CD57, and KLRG1, suggesting expansion of an antigen-experienced terminal effector memory population with potential for cytotoxicity in the context of MDS (**Extended Data Fig. 8a-b**).

**Fig. 4:**
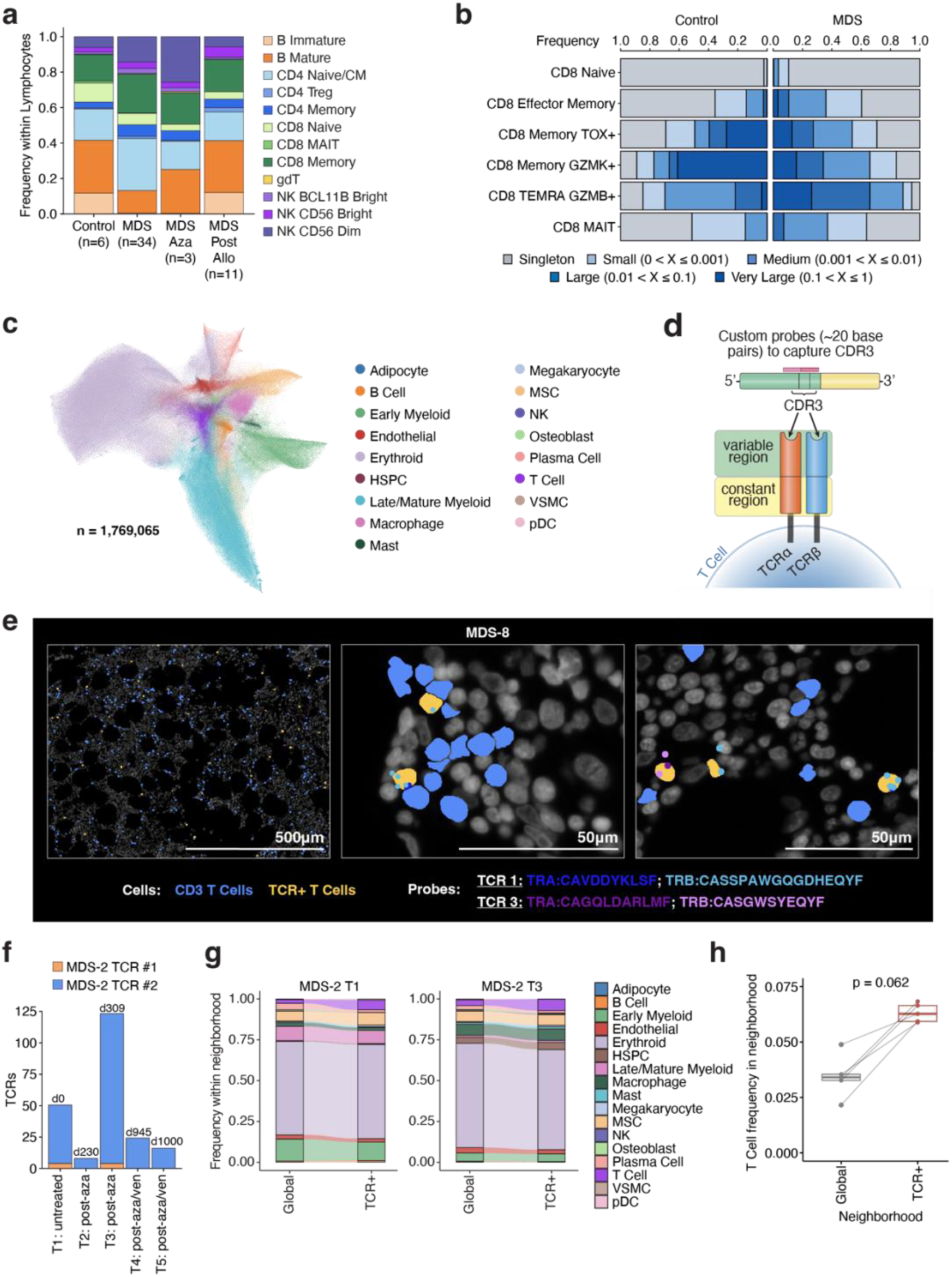
*In situ* spatial detection and localization of T cell receptors (TCRs) in human MDS bone marrow. **a**, Frequency of lymphocyte populations from bone marrow aspirates profiled by single cell CITE-seq across control (n=6), untreated MDS (n=34), azacitidine-exposed MDS patient samples (n=3), and MDS patients treated with allogenic hematopoietic stem cell transplantation (n=11). **b**, Clonal frequency of TCRs in CD8^+^ T cell subsets from control (left) and MDS (right) patient bone marrow aspirate samples, profiled by single cell TCR-seq, across a range of clone sizes from singletons to very large clones. **c**, UMAP representation of 1,769,065 single cells colored by distinct cell populations from Xenium spatial analysis of core bone marrow biopsies from control (n=3) and MDS (n=13) specimens. **d**, Schematic of custom probes to capture specific CDR3 sequences in TCR-alpha and TCR-beta. **e,** Representative images of T cells expressing defined TCRs detected in MDS-8 patient marrow. **f**, Bar plot of longitudinal spatial TCR tracking in MDS-2 across five timepoints depicting number of TCR^+^ cells recovered in each sample and colored by TCR. Timepoints in days are displayed above each bar. **g**, Alluvial plots comparing global cell type frequencies to cell type frequencies within 15 µm of a TCR^+^ cell in MDS-2 T1 (left) and T3 (right). **h,** Paired boxplots comparing the global frequency of T cells to the frequency of T cells within 15µm of a TCR^+^ cell in each sample with targeted TCRs. P-value was calculated by paired Wilcoxon signed-rank test.

Having defined the distinct localization, composition, and phenotype of T cells throughout the marrow of patients with MDS, we turned to single cell TCR analyses to quantify the diversity of the MDS T cell repertoire at diagnosis and over time. In both controls and patients with MDS, TCR repertoire clonality, a normalized metric ranging from 0 (all T cell clones are equally represented) to 1 (all T cells derived from a single clone) was greater in CD8 than CD4 T cells. Moreover, in MDS the most abundant CD8 T cell clones fell within the *GZMB^+^* CD8 terminal effector memory compartment; we did not identify abundant clonal Tregs in MDS or control samples (**Fig. 4b and Extended Data Fig. 9c-d**). Longitudinal assessment of the T cell repertoire across five MDS patients showed that dominant memory CD8 clones persisted over time prior to allogeneic transplantation; however, there was near complete replacement of T cell clones post-transplant (**Extended Data Fig. 9e**).

Having identified highly abundant memory CD8 T cell clones in patients with MDS, we sought to determine their *in situ* distribution. We designed probes for paired TCR-alpha and TCR-beta chains to localize individual patient-specific T cell clones within single T cells in the marrow. We selected the 14 most abundant TCR clones by frequency identified from four MDS patient aspirate samples and designed probes to spatially resolve TCRs within core biopsies. To maximize specificity, probes targeted the CDR3 region of both the TCR-alpha and TCR-beta chains, and these probes were included in a second targeted panel for spatial analysis of TCRs (**Fig. 4c-d, Extended Data Fig. 9f,** and **Supplementary Tables 4-6**). With this approach, we detected a total of 977 MDS patient- and T cell-specific TCR^+^ T cells within the marrow from three out of four MDS patients (see **Methods**, **Fig. 4e**, and **Extended Data Fig. 10a-e**), including in five longitudinal bone marrow biopsies from a patient over the course of MDS progression from diagnosis to 1,000 days after the initial bone marrow assessment (**Fig. 4f-g**, and **Extended Data Fig. 10f**). Cell types expressing the TCRs of interest from CITE-seq correlated with detectable TCR-specific cell types in the spatial data with the vast majority being memory CD8 T cells, validating our phenotypic definitions across modalities (**Extended Data Fig. 10g**). These highly abundant T cell clones were distributed throughout the marrow without spatial bias, reflected by their neighboring cell composition closely mirroring that of the global tissue composition. Compared to the global distribution of T cells, these TCR^+^ clones tended to be closer to other CD4 and CD8 T cells within the marrow (**Fig. 4h** and **Extended Data Fig. 10h-i**). Notably, only 3.2% of spatially resolved TCR^+^ T cells were located within immune aggregates (**Extended Data Fig. 10j**). These data align with our finding of increased memory CD8 T cells outside of immune aggregates (**Extended Data Fig. 5f**), some of which represent clonally expanded T cell subsets.

### *In situ* spatial detection of mutant cells in early stage MDS

One aspirational goal in the study of hematopoietic disorders originating in the bone marrow has been to delineate where malignant and disease-initiating cells are located within the marrow environment. Utilizing clinical targeted genomic DNA sequencing data obtained from our cohort of MDS patients, we designed probes to detect somatic single nucleotide variants (SNV), insertions, and deletions. We designed custom probes aimed at detecting specific mutations in *SF3B1, U2AF1, ASXL1, CUX1, IDH1*, and *DNMT3A* identified as somatic mutations in this MDS patient cohort based on clinically targeted bulk DNA sequencing data from aspirated bone marrow mononuclear cells (**Supplementary Table 5**). To improve specificity of mutant cell detection in MDS, we leveraged prior knowledge that somatic mutations in SF3B1 result in expression of aberrant RNA isoforms which uniquely and specifically track with mutant cells^32,33^. Spatial detection of SF3B1 mutant cells is particularly relevant in MDS given that SF3B1-mutant MDS represents a specific MDS subtype with unique therapeutic and prognostic relevance^34^. We incorporated custom probes targeting two previously defined SF3B1 mutation-specific aberrant RNA splicing events in *ORAI2*^35,36^ and *ZDHHC16* wherein the SF3B1 mutation promotes expression of intronic sequences not normally seen in SF3B1 wild-type cells^37,38^ (**Fig. 5a**). We thereby utilized SF3B1 mutant-specific alternative splicing events as a surrogate method for direct SF3B1 mutation identification (**Supplementary Table 5**). These probes were applied to bone marrow core biopsies from three control subjects and nine MDS patients, five of which were defined to have a mutation in at least one of these genes at >2% variant allele frequency from the clinical DNA sequencing data (**Supplementary Table 6**).

**Fig. 5:**
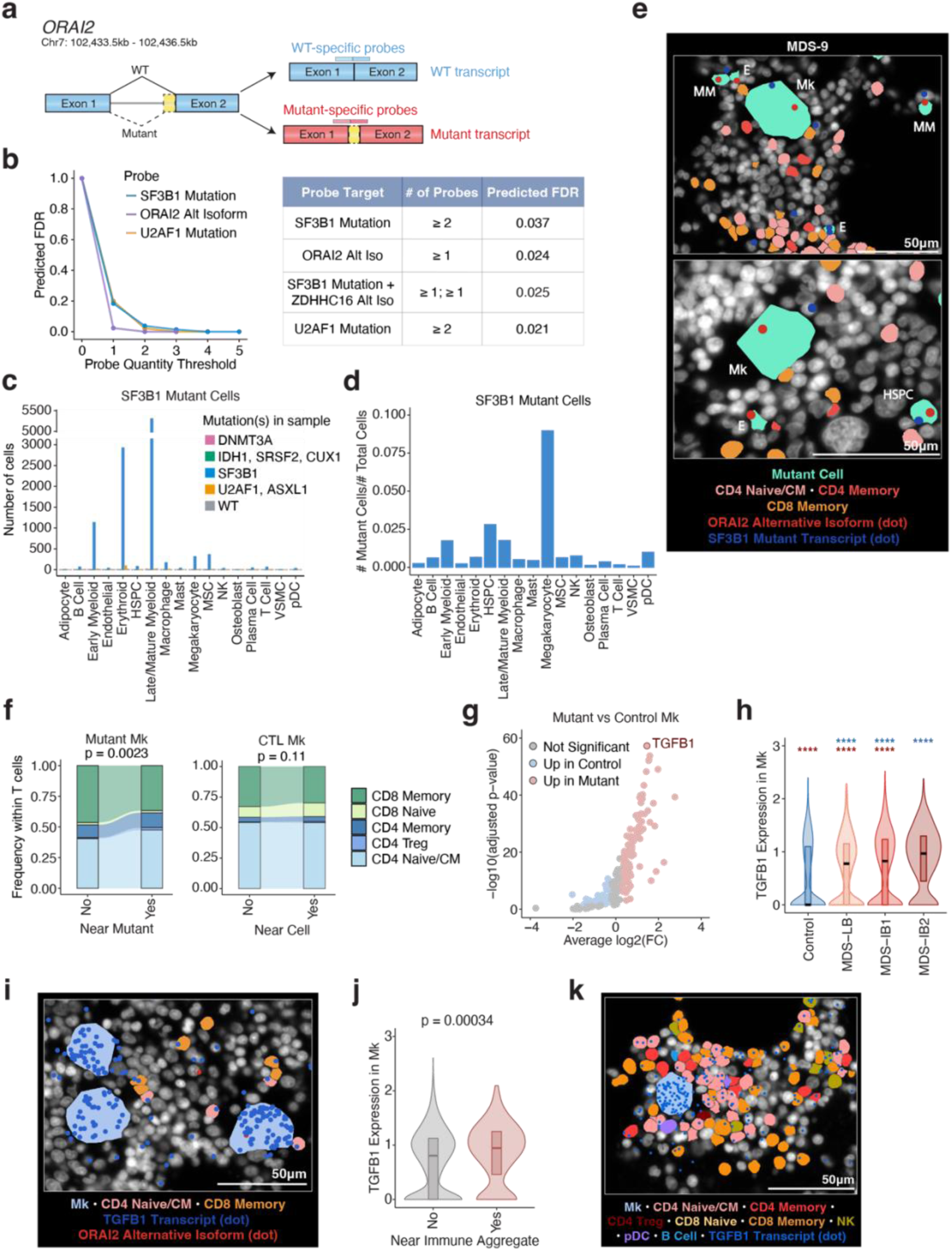
Spatially resolved mutant cells in human MDS bone marrow. **a**, Schematic of custom probes to detect normal and SF3B1-mutant induced aberrant *ORAI2* mRNA isoforms. **b,** Left, predicted false-discovery rate (FDR) of *in situ* somatic mutation and aberrant RNA isoform detection based on number of probes in cell. Right, table of probe, number of probes for assigning mutant cells, and corresponding FDR. **c,** Number of mutant SF3B1 cells as defined by the presence of SF3B1-mutant specific *ORAI2* aberrant RNA isoform, the combination of ≥1 *ZDHCC16* aberrant RNA isoform + ≥1 SF3B1-mutant probe, or ≥2 SF3B1 mutant probes in each cell type grouped by sample mutation status across 16 (n=3 control, n=13 MDS) patient samples. **d**, Proportion of SF3B1 mutant cells in each cell type across n=5 mutant SF3B1 MDS samples. **e,** Representative images of mutant cells (turquoise) with detection of SF3B1-specific aberrant *ORAI2* isoform (red dot) or *SF3B1* mutant transcript (blue dot) in SF3B1 mutant MDS patients. Mutant cell types are labeled in white. MM, mature myeloid; HSPC, hematopoietic stem and progenitor cell; E, erythroid; Mk, megakaryocyte. **f,** Left, alluvial plot comparing the frequency of T cell subtypes between T cells far (>15 µm) from a mutant Mk cell and T cell subtypes near (within 15 µm) a mutant Mk cell in SF3B1-mutant MDS samples. Right, alluvial plots comparing the frequency of T cell subtypes between T cells far (>15 µm) from a Mk cell and T cell subtypes near (within 15 µm) a Mk cell in control samples. P-values are calculated with a chi-squared test on T cell subtype counts between “No” and “Yes” groups. **g,** Volcano plot of differential gene expression between mutant megakaryocytes in SF3B1 mutant samples and megakaryocytes in control samples. **h**, Violin plots of *TGFB1* expression in megakaryocytes in control (n=13) and MDS (n=30) patient core biopsies regardless of megakaryocyte mutation status. At most, 100 megakaryocytes were randomly sampled from each sample. Asterisks denote p-values calculated from Wilcoxon rank-sum test comparing to control samples (blue) and MDS-IB1 samples (red), ****p ≤ 0.0001. Internal box represents interquartile range, and the line represents the median. **i,** Representative image of megakaryocytes with *TGFB1* transcript probes (blue dot) in a mutant SF3B1 sample. **j,** Violin plots of TGFB1 expression in megakaryocytes near (within 15 µm) and far (>15 µm) from an immune aggregate across MDS (n = 18) patient core biopsies regardless of megakaryocyte mutation status. At most, 50 megakaryocytes were randomly sampled from each sample for each group (“Yes”, “No”). P-value was calculated with a Wilcoxon rank-sum test. **k,** Representative image of a megakaryocyte expressing *TGFB1* transcript probes (blue dot) adjacent to an immune aggregate. Mk, megakaryocyte; NK, natural killer cell; Treg, regulatory T cell; pDC, plasmacytoid dendritic cell.

We examined the specificity of each mutation-targeting probe by comparing mutant probe expression between samples with and without each corresponding mutation. The specificity of probes was variable across mutations, with the highest specificity noted for the probe designed to detect SF3B1^K700E^ (c.2098A>G) and U2AF1^S34F^ (c.101C>T) mutations (**Fig. 5b** and **Extended Fig. 11a,b**). The lack of specificity of the other probes may be due to the limited change in nucleotide sequence imparted by these mutations or suboptimal expression of the mutant transcript. Detection of the *ORAI2* aberrant RNA isoform was highly specific for SF3B1 mutant cells (predicted false discovery rate (FDR) of 0.024). Defining mutant cells as having ≥1 *ORAI2* aberrant isoform transcript identified >8,000 mutant cells across five SF3B1 mutated patient samples (**Fig. 5b** and **Extended Data Fig. 11c,d**). Requiring ≥2 SF3B1 mutant probes or ≥1 SF3B1 mutant probe in combination with ≥1 *ZDHHC16* SF3B1-mutant specific isoforms (all methods with predicted FDR <5%) further refined *in situ* SF3B1 mutant detection (**Fig. 5b-d** and **Extended Data Fig. 11c-g, 12a-c**). These data provide proof-of-concept spatial detection of somatic mutant transcripts across key MDS cell populations and identification of hematopoietic clones *in situ* in human bone marrow.

### Increased TGFβ secretion by mutant megakaryocytes in MDS impairs local T cell cytotoxicity

Visualization of SF3B1 and U2AF1 mutant cells revealed mutant erythroid cells and megakaryocytes abutting CD4 and CD8 T cells across multiple patients (**Fig. 5e, Extended Data Fig. 11g, 12c**). Quantification of T cell subsets around mutant cells revealed that SF3B1-mutant erythroid and megakaryocytic cells each had distinct compositions of T cell neighbors, which was not observed in control marrows. Specifically, we observed a relative decrease in CD8 memory T cell frequency among T cells neighboring SF3B1-mutant megakaryocytes (**Fig. 5f, Extended Data Fig. 12d-e**).

Given this spatial reorganization of T cells around mutant cells, we performed differential gene expression analysis comparing mutant cell types to controls, an analysis that has previously been limited in megakaryocytes due to challenges in capturing primary megakaryocytes in single cell suspensions. Strikingly, the top differentially expressed gene in SF3B1-mutant megakaryocytes compared to control megakaryocytes was *TGFB1* which encodes TGFβ, a molecule known for its role in suppression of anti-tumor immunity across malignancies^39^ (**Extended Data Fig. 12f**). Furthermore, there was significantly increased expression of TGFβ in MDS megakaryocytes compared to controls (n=13) at all stages of MDS (n=30) (**Fig. 5g)**. Notably, the highest TGFβ expression was observed in patients with MDS-IB2 (**Fig. 5h-i).** Moreover, we found higher TGFβ expression in megakaryocytes near immune aggregates in MDS and near MSCs, suggesting a potential role for spatially restricted megakaryocyte-derived TGFβ signaling in shaping the local immune microenvironment (**Fig. 5j-k, Extended Data Fig. 12g**). Of note, mutant myeloid cells expressed distinct immunosuppressive molecules amongst SF3B1 mutant cells such as VISTA and galectin-9, suggesting distinct mechanisms of immune evasion based on mutant cell types (**Extended Data Fig. 12h**).

Across many cancers, TGFβ plays a role in limiting T cell cytotoxicity by dampening CD8 T cell effector function and reducing expression of cytolytic molecules^40^. We therefore asked whether the high TGFβ expression found in MDS patient marrow might reshape the profile of MDS T cells. Incubation of TGFβ with primary MDS mononuclear cells revealed dose-dependent suppression of perforin and granzyme B expression on CD8 T cells *in vitro*, supporting a TGFβ-mediated inhibitory effect on CD8 T cell markers of cytotoxicity (**Fig. 6a**). To validate these findings and assess their biological relevance in the MDS bone marrow, we examined our spatial transcriptomics data to compare the gene expression profiles of CD8 memory T cells based on their proximity to mutant megakaryocytes. Notably, we observed significant downregulation of cytotoxicity-related genes, including perforin (*PRF1*) and Tbet (*TBX21*), in CD8 T cells physically adjacent (≤20µm distance) to mutant megakaryocytes, compared to CD8 memory T cells further (>50µm) from mutant megakaryocytes (**Fig. 6b**). Together, these data suggest a model whereby increased TGFβ secretion by mutant megakaryocytes in MDS reshapes the immune microenvironment through suppression of local CD8 T cell immunity, potentially contributing to immune evasion in MDS.

**Fig. 6:**
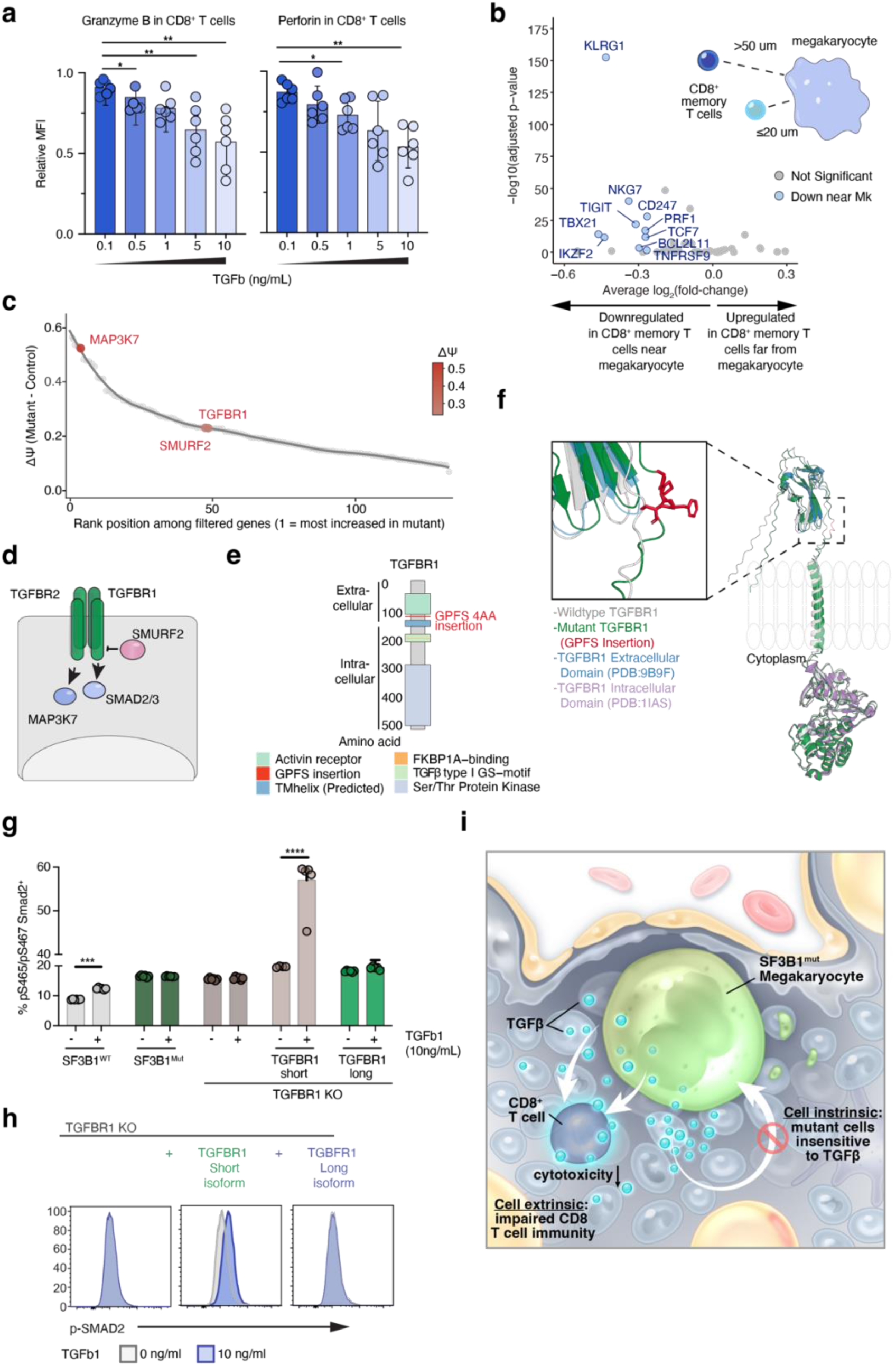
Impaired TGFβ signaling by SF3B1 mutant cells in MDS suppresses local T cell immunity in the bone marrow. **a,** Bar plots of relative median fluorescence intensity (MFI) of granzyme B and perforin from intracellular cytokine staining of primary untreated MDS CD8 T cells after exposure to increasing doses of TGFβ. Relative MFI values are shown as mean ± SD. Data from n=6 MDS patients. Mann-Whitney U test used for statistical analysis (*p<0.05, **p<0.01). **b,** Top, volcano plot of differentially expressed genes in CD8 memory T cells close to (≤20µm) versus far from (>50µm) megakaryocytes. Bottom, schematic of CD8 memory T cells based on their distance to nearest megakaryocytes from spatial data. **c,** Relative percent spliced in (dPSI) value of differential 3’ splice site in mRNAs from SF3B1 mutant MDS patient bulk RNA-seq versus SF3B1 wild-type MDS. Highlighted names indicate mRNAs encoding proteins involved in TGFβ signaling. **d,** Diagram of proteins (in green) involved in TGFβ signaling which undergo aberrant RNA splicing in SF3B1 mutant MDS. **e,** Protein diagram of TGFBR1 with red indicating insertion of four amino acids (GPFS) encoded by the long mRNA isoform promoted in SF3B1 mutant cells. **f,** Alpha fold model of mutant TGFBR1 (green) overlaid with published crystal structure of wild-type TGFBR1 (gray). The GPFS amino acid insertion seen in SF3B1 mutant cells is indicated in red in the inset. **g,** Percentage (%) of phospho-SMAD2 Serine 465/467 (pS465/467) in K562 cells with the indicated genetic alterations in SF3B1 or TGFBR1. Mean ± SD. Two-way ANOVA. ***p<0.001, ****p<0.0001. **h**, Representative flow cytometry histograms of p-SMAD2 S465/467 from (g). **i,** Schematic of an SF3B1-mutant megakaryocyte with increased TGFβ production suppressing cytotoxic activity of nearby CD8^+^ T cells (cell-extrinsic effect); mutant cell simultaneously exhibits impaired TGFβ sensing (cell-intrinsic effect).

### Aberrant TGFβ signaling in SF3B1 mutant MDS

Due to the known clinical connection between TGFβ signaling suppression and therapeutic impact on SF3B1 mutant MDS specifically^41^ we set out to understand the impact of aberrant TGFβ signaling on SF3B1 mutant cells. Mutations in SF3B1 cause differential 3’ splice site usage leading to widespread production of mis-spliced mRNAs across many different pathways that drive cancer pathogenesis^32,37^. RNA-seq analysis of publicly available SF3B1 mutant leukemias revealed enrichment for differential 3’ splice site usage in mRNAs encoding a number of TGFβ signaling molecules including TGFBR1, a transmembrane receptor critical for TGFβ signaling, MAP3K7, a kinase downstream of TGFβ signaling and a number of cell death receptors, and SMURF2, a negative regulator of TGFBR1 protein abundance (**Fig. 6c-d** and **Extended Data Fig. 13a)**. Prior work from our group and others has revealed that SF3B1 mutations promote expression of an out-of-frame transcript in MAP3K7 resulting in downregulation of MAP3K7 mRNA and protein^37,42^. However, here we identify that mutant SF3B1 promotes use of a frame-preserving splicing event in TGFBR1 resulting in inclusion of four amino acids in the extracellular domain just distal to its activin binding domain (**Fig. 6e-f** and **Extended Data Fig. 13b**). These SF3B1 mutant-specific events were validated by RT-PCR in SF3B1^K700E^ knockin K562 isogenic cell lines as well as primary MDS patient bone marrow aspirate samples (**Extended Data Fig. 13c-e**).

The function of this specific isoform of TGFBR1 has not previously been evaluated in myeloid malignancies. We therefore assessed the impact of the SF3B1^K700E^ mutation versus TGFBR1 knockout (KO) within isogenic K562 cells. In addition, we included cells with isolated expression of the long versus short isoforms of TGFBR1 (expressed as cDNAs in the TGFBR1 KO SF3B1 wild-type cells) and confirmed that both long and short TGFBR1 isoforms traffic to the cell membrane (**Extended Data Fig. 13 f-g**). We found that SF3B1 mutant cells as well as cells with TGFBR1 deletion or isolated expression of the mutant-promoted TGFBR1 long-isoform have impaired signaling in response to TGFβ (**Fig. 6g-h**). Together, these data support a model whereby SF3B1 mutant cells have enhanced TGFβ secretion, resulting in local suppression of anti-tumor immunity, while simultaneously harboring cell-intrinsic alterations in sensing TGFβ (**Fig. 6i**).

## DISCUSSION

Extensive prior studies in mice, other model organisms, and primary human tissue have shown that the bone marrow is spatially organized into distinct niches, enabling cellular interactions between hematopoietic and non-hematopoietic cells that are critical to hematopoiesis. The advent of spatial imaging technologies has recently yielded several landmark high-resolution studies of the topography of primary human bone marrow *in situ* in disease-free samples^16,17^. To date, however, there have been limited single cell spatial transcriptomic studies to define molecular features of intact primary human bone marrow in the setting of MDS. As a result, the ecological structures and interactions that underlie progression have not been quantified.

Here we performed a comprehensive molecular analysis of human bone marrow *in situ* and spatially identified both hematopoietic and non-hematopoietic cell types, as well as TCRs, disease defining somatic mutations, and aberrant RNA isoforms. This was enabled by the usage of EDTA decalcification of fixed clinical bone marrow specimens from patients to allow for preservation of nucleic acids and downstream molecular analysis of intracellular RNA in whole tissues.

We applied custom targeted spatial gene expression panels to study human bone marrow core biopsies from patients with MDS. To confidently identify mutant cells, we designed custom probes targeting SF3B1 and U2AF1 somatic hotspot mutations, as well as SF3B1 mutation-derived aberrant RNA isoforms. This approach allowed us to capture multiple cell types, including those which are difficult to aspirate, such as megakaryocytes, endothelial cells, adipocytes, and stromal populations. Evaluation of the molecular characteristics of these cells in a spatial context is particularly relevant as non-hematopoietic cells within the marrow can functionally contribute to the aberrant clonal hematopoiesis characteristic of MDS^9,21,43^. We therefore prioritized our analyses on early-stage newly diagnosed or untreated MDS to capture the initial remodeling of the microenvironment prior to disease progression.

To enhance our ability to define and analyze biologically relevant structures in bone marrow, a tissue that lacks typical spatial landmarks, we developed two computational methods called SAND and DUNE to disentangle the microscopic ecological determinants of disease. This analysis revealed that human early-stage MDS bone marrow is characterized by abnormal cell-cell relationships and distributions of cell types. SAND enabled refinement of megakaryocyte cell segmentation as well as detection and characterization of several key marrow structures including cytokine clusters and immune aggregates with several features similar to tertiary lymphoid structures which have not been captured in prior studies of the bone marrow at this level of detail. Importantly, this tool can be applied to other heterogenous and morphologically unstructured tissues allowing for the identification of discrete structures for broad computational spatial analyses. DUNE revealed that MDS marrow is stratified from normal marrow along two orthogonal niche axes, an erythroid axis enriched in earlier stage disease and a myeloid axis enriched in later stage disease. This result suggests the tantalizing possibility that spatial information uniquely classifies patient disease state, with potential implications for clinical decision-making.

The spatially informed genotyping of SF3B1 mutant cells revealed a TGFβ-mediated interaction by which SF3B1 mutant megakaryocytes suppress the functionality of abutting T cells. We extensively validated this interaction, confirming dose-dependent TGFβ-mediated suppression of MDS patient T cells *in vitro*. At the same time, we identified a novel TGFβ splice variant promoted by the SF3B1 mutation which blunts sensing of TGFβ in mutant cells (and this occurs in the context of additional mis-splicing events perturbing intracellular TGFβ signaling). The SF3B1 mutant promoted isoform of TGFBR1 may attenuate TGFβ signaling by negatively impacting its heterodimerization with TGFBR2 and/or affecting post-translational modification of TGFBR1 which reduce its ability to transmit signaling. This finding has several important implications. First, these data suggest a model whereby the mutant cells may be clonally selected through a combination of TGFβ-mediated suppression of local tumor immunity while allowing the mutant cells to persist in the setting of high TGFβ. Second, while pharmacologic suppression of TGFβ signaling is known to promote terminal red blood cell differentiation in SF3B1-mutant MDS, it will also be interesting to evaluate the impact of these agents on the bone marrow immune microenvironment. Finally, it is important to note that the SF3B1 mutant promoted TGFBR1 has a novel extracellular domain, suggesting the possibility of therapeutic targeting of mutant induced cell surface epitopes such as this in the future.

Inflammation has long been suggested to drive dysfunction in the hematopoietic niche in MDS patients^10,44^, and there are established immunosuppressive therapies to improve cytopenias in a subset of low-risk MDS^45,46^. These findings have motivated multiple therapeutic intervention studies in MDS patients including agents targeting IL-1, TGFβ, and IRAK1/4 signaling amongst others^10^. Here we identify that the bone marrow in patients with MDS contains immune aggregates with distinct cellular and molecular composition relative to the bone marrow of age-matched control subjects, including increased *TCF7* expression. *TCF7* encodes a transcription factor defining memory T cells with stem-like properties, which serve as an “effector T cell reserve” in the context of chronic inflammation and antitumor immunity^47^. These data provide evidence for altered adaptive immune response in early stage MDS and SF3B1-mutant, favorable-risk disease. Recent discovery of shared neoantigens in splicing factor mutant cancers^48,49^, combined with the increased frequency of stem-like memory T cells in SF3B1 mutant MDS specifically here suggests an intriguing possibility of a neoantigen-specific immune response in this subset of MDS. Future efforts to spatially evaluate antigen-specific T cells identified in SF3B1 mutant MDS will be helpful to further evaluate this potential connection.

The spatial localization of individual T cell clones defined by their TCR-alpha and or - beta chain represents a technical advance with broad implications for analysis of tissue-based pathology. The single cell resolution of our approach enables identification of T cell clones based on paired receptor chains and their surrounding cellular compartments *in situ*. This method can be applied to fixed tissues, which is distinct from previous methods which require fresh frozen tissue^50,51^. At the same time, the present method relies on targeted sequencing to identify specific TCR transcripts. As such, the multiplexing and scalability of our approach is more limited than previously described whole transcriptome approaches which allow for capture of TCR clones (including spatial VDJ^51^ and Slide-TCR-seq^50^) without prior knowledge of TCR sequence, albeit at the cost of single cell resolution.

One challenge in the clinical assessment and study of MDS is that the bone marrow consists of an admixture of cells with disease-defining somatic mutations interspersed with wild-type HSPCs and mature hematopoietic cells. Prior studies have attempted to address this challenge by genotyping of cellular material dissected from known geographic regions of bone marrow^52^. While these studies made important insights into localization of mutant cells, such an approach lacks fine-scale information about the spatial context of mutant cells and surrounding non-mutant cells. The ability to now genotype clonal mutations *in situ* in combination with capturing gene expression within mutant cells and their ecological milieu is likely to introduce new perspectives into how cancer is initiated and maintained. Furthermore, there is hope that understanding the interactions between mutant cells and their physically associated non-mutant cells will highlight new therapeutic interventions to treat cancer by perturbing these interactions.

## MATERIALS AND METHODS

### Patient Samples and Approval

Studies were approved by the Institutional Review Boards (IRB) of Memorial Sloan Kettering Cancer Center and conducted in accordance with the Declaration of Helsinki protocol. Patient samples were collected and analyzed under MSK IRB protocols (#06-107, #x19-016, #12-245, #16-834). Primary bone marrow core biopsies and bone marrow mononuclear samples from de-identified patients with MDS or control subjects were utilized for Xenium spatial analysis, and/or single cell CITE- and TCR-seq, and/or H&E staining, and/or multiplexed immunofluorescence. Bulk mutational genotyping of each sample was performed in the clinical environment by the MSKCC IMPACT assay as previously described^53^.

### Specimen Collection and Processing

A list of patients which meet criteria for a diagnosis of MDS based on WHO 2022 classification criteria was curated based on review of clinical information, peripheral blood counts, bone marrow morphologic review, and supporting tandem cytogenetic and molecular findings. Specimens that met criteria for lymphoid clonal bone marrow disorders were excluded. Control bone marrow core biopsies were collected from healthy bone marrow donors (n=5) and lymphoma negative staging marrows (n=10). Post-transplant bone marrow core biopsies (n=5) were collected from patients with a history of MDS who had been in complete remission for ≥ 722 days (range 722-1242 days) post allogeneic hematopoietic stem cell transplant with undetectable minimal residual disease, assessed by both multiparametric flow cytometry and next generation sequencing (details in **Supplementary Table 2**). Bone marrow core biopsies were either prospectively collected or retrieved from the archive of the Pathology Department of MSKCC. In both instances, adequacy of bone marrow spaces in sampled core biopsies for all downstream studies was evaluated via H&E review by a board-certified hematopathologist. Bone marrow core biopsies were fixed in 10% neutral buffered formalin overnight. After rinsing with running cold water for 5 minutes, cores were placed in cassettes in Milestone Mol-Decalcifier solution (unbuffered 10% EDTA decalcification solution) and run through an automated decalcification protocol (Milestone KOS instrument, 37 °C – 45 °C; EDTA, pH 7.2) for 7 to 10 hours. After rinsing with cold water (2-5 minutes), cassettes were placed into tissue processors followed by tissue embedding into paraffin blocks. Formalin-fixed, paraffin-embedded (FFPE) cores were sectioned with a microtome (Leica Biosystems) and 5μm sections were either directly mounted on a specialized Xenium slide (10x Genomics) for spatial imaging or conventional glass slides (Fisherbrand, Superfrost Plus) for use in validation studies with immunofluorescence microscopy.

### Xenium Run and Post-Xenium Tissue and Image Processing

Two pre-designed RNA Xenium panels were paired with custom probes and used for spatial analysis. Panel 1.0 (61 total patients; 15 controls, 41 MDS and 5 post-transplant) consisted of 377 genes from the Human Multi-Tissue and Cancer pre-designed panel (10x Genomics) combined with custom designed probe panels (A-C) targeting 56-100 genes additional genes (**Supplementary Table 1**). Xenium panel 2.0 (12 total patients; 3 controls, 9 MDS) consisted of probes targeting 380 genes from the Immuno-Oncology pre-designed panel (10x Genomics) combined with custom designed probe panel D targeting 100 additional genes including SNVs, splice isoforms, and TCRs (**Supplementary Tables 4-6**). Xenium experiments were performed according to the manufacturer’s directions.

Following a Xenium Analyzer (10x Genomics) imaging run, Xenium slides were immersed in Quencher removal solution, followed by a modified H&E staining protocol (protocol available on 10x Genomics website). Subsequently, H&E stained Xenium slides were scanned via a Pannoramic scanner (3DHistech, Budapest, Hungary) using a 20x/0.8NA objective. A hematoxylin channel was color deconvolved from the whole slide image of H&E-stained tissue. This channel was aligned to the DAPI channel from Xenium via linear registration (imregcorr function in MATLAB’s Image Processing Toolbox) followed by nonlinear registration (imregdemons function in MATLAB’s Image Processing Toolbox). The resulting calculated transformation was applied to the whole slide image of H&E-stained tissue, yielding an H&E image in the same coordinate space as the Xenium image. Code for Xenium-H&E image alignment will be made publicly available upon peer-reviewed publication. Post-Xenium H&E images were also aligned utilizing Image Alignment software in Xenium Explorer 4.0 (10x Genomics).

### Xenium Preprocessing

Spatial transcriptomics data was first segmented into individual cells. When available, the Xenium multimodal cell segmentation was utilized. Otherwise, cell segmentation was performed using Baysor v0.6.2 with cells defined by nuclear transcripts determined by Xenium Analyzer as a prior cell segmentation. Additionally, the following parameters were used: 10 minimum molecules per cell, 15 molecule clusters, and a prior segmentation confidence of 0.8. Xenium Ranger v2.0.1 was used to import Baysor segmentation outputs.

Data quality control steps included selecting for cell-dense regions corresponding to intact marrow spaces and excluding regions of blood clot and aspirated marrow spaces, as well as filtering cells by quantity of genes expressed, transcript count, and presence of nuclear transcripts resulting in a population of high-quality cells. Specifically, cell feature matrices from Xenium Ranger were imported into Scanpy v1.9.1 as AnnData objects (anndata v0.9.2). Cells within sparse regions (sum of distances to 50 nearest neighbors >2^12^), cells expressing less than 5 genes, cells with low transcript counts (<5 to 8 transcripts depending on distribution within sample), and cells with no nuclear transcripts were removed. Transcript counts were normalized by total transcripts per cell to a target sum of 50. Sample AnnData objects were combined and visualized using UMAP which was derived from first computing a nearest neighbor graph (15 neighbors, top 20 PCs), Leiden clusters (resolution=1.0), and a Partition-based Graph Abstraction (PAGA). The UMAP was then generated using the PAGA graph as the initial position and a minimum distance of 0.1. Low quality samples determined by low transcript counts per cell and clustering by sample in the UMAP were discarded.

### Xenium Cell Typing

Cells were manually typed based on Leiden clustering and differentially expressed genes in consultation with a board-certified hematopathologist.

### SAND (Spatial Adjacency Network Detection)

For identifying 2-D spatial clusters, SAND accepts a matrix of coordinates (x,y) along with annotations (cell type, gene name, etc). SAND first creates a neighborhood graph using the coordinates which can either be KNN-based (default) or radius-based. The KNN graph (k = number of nearest neighbors) is further filtered such that nodes not belonging to feature categories of interest and edges with distances greater than or equal to an optional specified cutoff (r) are removed. Connected components in the filtered graph are identified with union-find and outputted. SAND was used for refining megakaryocyte segmentation (r = 5, see below), identifying immune aggregates (k = 10, r = 15), and identifying cytokine/chemokine clusters (r = 10, with a gene filtered transcript matrix input). Immune aggregates from SAND were further filtered such that each aggregate contained at least 10 T cells and 20% of the aggregate was composed of T cells. This filtering criteria was determined to exclude “aggregates” driven primarily by an abundance of B cells in rare samples. Transcript clusters were filtered such that each cluster contained at least 10 transcripts. Code for SAND will be made publicly available upon peer-reviewed publication.

### Megakaryocyte Segmentation

To merge over-segmented megakaryocytes, a convex hull was computed over all vertices corresponding to cells within each connected component utilizing the “convex_hull” function from shapely v2.0.6. The resulting polygon defined the new boundary of the re-segmented megakaryocyte, and its centroid was re-computed accordingly. Gene expression counts were aggregated by summing counts across all constituent over-segmented cells within each connected component. These re-segmented megakaryocytes were then re-integrated into the original AnnData object with new centroids, replacing all previously over-segmented megakaryocyte cells. Given the large size of megakaryocytes, radius-based spatial analyses of megakaryocytes are computed from polygon vertices rather than the centroid (**Fig. 3e, 4g, 5f, 5j, 6b** and **Extended Data Fig. 3g, 10i, 12g**). For visualization in Xenium Explorer, the Xenium Ranger v4.0.0 “import-segmentation” function was used to import the new megakaryocyte segmentation from cell polygons.

To re-segment over-segmented megakaryocytes, we applied SAND to vertices of cell polygons initially annotated as megakaryocytes. One sample (region1_0069146), characterized by a high abundance of micro-megakaryocytes, was excluded from this step, and the original Xenium multimodal segmentation was retained. Vertices located within 5 µm of one another were considered neighbors, enabling the identification of spatially neighboring, over-segmented cells for merging. Specifically, we constructed a radius-based neighbor graph (r = 5µm) over polygon vertices and identified connected components with SAND which were then mapped back to their corresponding cells. Components sharing cells were further consolidated using union-find. Each resulting connected component therefore represented a set of over-segmented megakaryocytes to be merged into one.

### DUNE (Discovery by Unsupervised Niche Enrichment)

To compare the spatial architecture between individual bone marrow samples, we propose a method for unsupervised analysis of cellular niches. First, we introduce a framework to quantify the enrichment of cell niches which will enable the identification of over- and under- represented local structures. For each cell, we define its niche as the cell type count vector over its *k* nearest neighbors together with the cell itself (**Fig. 2A**). For a given niche **n**, let

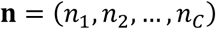

denote the counts of each of *C* cell types of **n**, such that

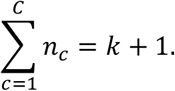

For a given sample *S*, let

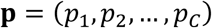

denote the global proportions of each cell type, where *p_c_* is the proportion of cell type *c* in *S*. Under a random mixing model with replacement, the likelihood of observing a niche **n** in *S* is given by a multinomial distribution:

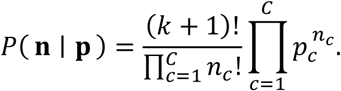

Let *N* be the total number of cells (and therefore individual niches) in sample *S*, and let *N(**n**)* denote the number of occurrences of niche **n**. The observed proportion of this niche is:

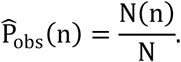

With this, we define the niche enrichment score as the ratio of the observed proportion to its expected likelihood under the multinomial model:

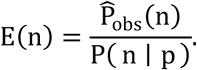

To perform unsupervised niche enrichment analysis, we first filter niches by entropy and variance of their enrichment scores. Entropy-based filtering is performed to remove features that are dominated by rare outlier samples and is achieved by computing the Shannon entropy for each niche. Specifically, for a given niche, enrichment scores (log_2_ transformed and z-score scaled) across samples are first discretized using quantile binning where values are partitioned into n = 4 bins. To ensure unique bin boundaries, duplicate quantile edges are removed. If there are fewer than two remaining bins, the entropy is set to 0. Otherwise, each enrichment value is then assigned to a bin, and the Shannon entropy is computed as:

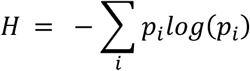

Where *p_i_* is the probability of observations in bin *i.* For principal component analysis and unsupervised clustering, we filter out niches with entropy = 0 and variance < 1 (where variance is calculated on log_2_ transformed enrichment scores). Code will be made publicly available upon peer-reviewed publication.

The matrix of niche enrichment scores was loaded into an AnnData object, and scores were log transformed. Following this, scores corresponding to niches with less than three occurrences within a sample were masked with 0. Finally, scores were z-score normalized across samples. The “.var[“highly_variable”]” field of the Anndata object was set to niches with entropy > 0 and variance ≥ 1. Following this, principal component analysis was performed utilizing the Scanpy “pca” function, and Leiden clusters were computed by first constructing a neighborhood graph with the “neighbors” function (n_neighbors = 5, n_pcs = 15) followed by “leiden” (resolution = 1). Differential niche enrichment analysis was computed with the “rank_genes_groups” function (method = “wilcoxon”). Niches with adjusted p-values < 0.05 were determined as “differentially enriched”. Sparse regions were excluded from analysis (region2_0042591, region2_0078981, region1_0078989, region4_0065604).

### Pairwise Neighborhood Enrichment Analysis

Neighborhood enrichment scores between each pair of cell types in each sample were calculated using the “nhood_enrichment” function in Squidpy v1.2.3 (for details see Neighborhood Enrichment Test in Methods section^22^) with 5 nearest neighbors which is a permutation-based method of determining neighborhood enrichment. Sparse regions were excluded from analysis (region2_0042591, region2_0078981, region1_0078989, region4_0065604).

### Multimodal Single Cell Sequencing

Previously isolated and viably frozen bone marrow mononuclear cells (BMNCs) from patient samples and controls were thawed and introduced to prewarmed and filtered 1640 RPMI (Gibco, catalog #11875093) with 10% FBS. Cells were suspended in Human FcR Blocking Reagent (Miltenyi Biotec, catalog #130-059-901), diluted in sterile-filtered DPBS (Corning, catalog #21-031-CM) + 0.5% BSA (Fisher, catalog #F2442-500ML), and incubated for 10 minutes at 4°C. Brilliant Staining Buffer (BD), TotalSeq^TM^ Oligonucleotide anti-Human Hashtag antibodies (Biolegend; C0251 catalog # 394661, C0252 catalog # 394663, C0253 catalog # 394665, C0254 catalog # 394667, or C0258 catalog # 394675), surface antibody mix (CD3 PE-Cy7, Invitrogen; CD45 BV510, Biolegend; CD34 APC, BD Biosciences; was added to the FcR Blocking solution and incubated at RT for 30 minutes in the dark. Cells were washed with filtered DPBS + 0.5% BSA, filtered using 30 µm filters (Sysmex), and resuspended in filtered DPBS + 2% BSA + DAPI (25 ng/sample). CD45dim and CD45+ cell populations were purified using Fluorescent Activated Cell Sorting (FACS), using the Aria3 Instrument (BD Biosciences). See (**Extended Data Fig. 14)** for gating strategy for sorting of CD45dim and CD45+ populations. Upon purification and collection of cell populations in Protein Lo-Bind Eppendorf Tubes #30108442, samples were pooled together based off cell number (with a goal of 50-100k cells per sample) and washed with filtered Cell Staining Buffer (Biolegend, catalog # 420201). Cells were Fc Blocked using TruStain FcX (Biolegend) at 4°C for 10 minutes and then stained with Totalseq^TM^ Universal Cocktail (Biolegned, catalog #399905) for 30 minutes at 4°C. The cells were washed once with Cell Staining Buffer and twice with DPBS + 0.04% BSA and then submitted for sequencing.

Samples were subject to 10x Genomics 5’ single cell RNA sequencing combined with TCR sequencing and CITE-sequencing after sample hashing^54^. FACS-sorted cell suspensions were subject to the Chromium Single Cell instrument (10X Genomics) following the user guide manual for 5′ v2 chemistry. In brief, FACS-sorted cells were washed once with PBS containing 0.04% bovine serum albumin (BSA) and resuspended in PBS containing 0.04% BSA to a final concentration of 700–1,200 cells per μl. The viability of cells was above 80%, as confirmed with 0.2% (w/v) Trypan Blue staining (Countess II). Cells were captured in droplets. Following reverse transcription and cell barcoding in droplets, emulsions were broken and cDNA purified using Dynabeads MyOne SILANE followed by PCR amplification per manual instruction.

Approximately 10,000 cells were targeted for each sample (or as many as were available), and several samples were multiplexed together on one lane of 10X Chromium to reach 30,000 cells targeted (when 3 samples were pooled together), following cell hashing protocol^54^. The scRNA-seq and scTCR-seq libraries were prepared using the 10x Single Cell Immune Profiling Solution Kit, according to the manufacturer’s instructions. Briefly, amplified cDNA was used for both 5′ gene expression library construction and TCR enrichment. For gene expression library construction, amplified cDNA was fragmented and end-repaired, double-sided size-selected with SPRIselect beads, PCR-amplified with sample indexing primers (98 °C for 45 s; 14–16 cycles of 98 °C for 20 s, 54 °C for 30 s, 72 °C for 20 s; 72 °C for 1 min), and double-sided size-selected with SPRIselect beads. For TCR library construction, TCR transcripts were enriched from 2 μl of amplified cDNA by PCR (primer sets 1 and 2: 98 °C for 45 s; 10 cycles of 98 °C for 20 s, 67 °C for 30 s, 72 °C for 1 min; 72 °C for 1 min). Following TCR enrichment, the enriched PCR product was fragmented and end-repaired, size-selected with SPRIselect beads, PCR-amplified with sample-indexing primers (98 °C for 45 s; 9 cycles of 98 °C for 20 s, 54 °C for 30 s, 72 °C for 20 s; 72 °C for 1 min), and size-selected with SPRIselect beads. Final libraries (GEX, TCR and HTO) were sequenced on Illumina NovaSeq S4 platform (R1 – 26 cycles, i7 – 10 cycles, i5 – 10 cycles, R2 – 90 cycles).

### CITE-seq/TCR-seq Preprocessing

Cell Ranger filtered cell feature matrix outputs were imported as AnnData objects for preprocessing with Scanpy. Cells were initially filtered such that cells with less than 200 expressed genes were removed, and genes were filtered such that genes expressed in less than 10 cells were removed. Next, cells were filtered based on total counts excluding ribosomal and mitochondrial genes. Cells with less than 2^9^ to 2^10.5^ non-ribosomal/mitochondrial counts were removed, with the exact cutoff determined on a per sample basis by visualization of count density. Cells were then filtered such that those with greater than 5 to 8 percent mitochondrial content were removed (per sample basis). Doublet Detection v4.2 was used to annotate doublets. Leiden clusters (1.5 resolution) were computed for each sample, and clusters with greater than 30% doublet cells were removed. Gene expression was normalized by total counts per cell and log transformed (natural logarithm). Centered log ratio (CLR) transformation was applied to ADT raw counts for normalization using muon v0.1.6.

Samples were then combined into a single object and visualized with a weighted nearest neighbor (WNN) UMAP. The WNN UMAP was constructed by first computing highly variable genes from the RNA data (scanpy.pp.highly_variable_genes with default parameters) to identify the most informative features and applying PCA to both RNA and ADT (scanpy.tl.pca with default parameters). Next, to integrate the transcriptomic and proteomic data, the pyWNN package was utilized to compute a WNN graph. The number of PCs for RNA and ADT was set to 25 and 15 respectively, and a neighborhood size of 15 was used for WNN calculation. Leiden clustering with a resolution parameter of 1.5 was then performed using the WNN graph as the neighborhood structure (scanpy.tl.leiden). Subsequently PAGA analysis was performed (scanpy.tl.paga), and UMAP embedding was computed using the PAGA graph abstraction as the initial position and the WNN graph as the neighborhood structure (scanpy.tl.umap). TCR sequencing data was imported from CellRanger outputs with custom scripts. TCRs were annotated based on the beta chain. In the case where a cell contained multiple beta chains, the most frequent beta chain in the sample was used. Clonality was calculated as previously described^55^.

### CITE Cell Typing

Cells were manually typed based on Leiden clustering and differentially expressed genes in consultation with a board-certified hematopathologist. In some cases (for T cell subtypes), batch correction with harmonypy v0.0.9 was applied before clustering.

### Immunofluorescence Microscopy

Tissue sections were stained using a six-marker multiplex immunofluorescence (mIF) assay targeting MPO, CD20, CD3, CD45RO, TCF1, and CD8. Fluorophores were assigned as follows: Opal 480 (MPO), Opal 520 (CD20), Opal 570 (CD3), Opal 620 (CD45RO), Opal 690 (TCF1), and Opal 780 (CD8). Staining was performed on the Leica BOND RX automated platform (Leica Biosystems). Heat-induced epitope retrieval (HIER) was conducted using Epitope Retrieval Solution 2 (ER2, Leica) for 30 minutes. Each primary antibody—MPO (RM407, SAB56000306, Millipore Sigma), CD20 (E7B7T, 48750, Cell Signaling Technology), CD3 (LN10, NCL-L-CD3-565, Leica Biosystems), CD45RO (UCHL1, 55618, Cell Signaling Technology), TCF1/TCF7 (C63D9, 2203, Cell Signaling Technology) and CD8 (NCL-L-CB8-4B11, Leica Biosystems) — were applied sequentially for 30 minutes at room temperature, except for CD45RO, which was incubated for 60 mins. Antibody dilutions were optimized through prior titration on normal and neoplastic control tissues. Secondary detection employed species-specific SuperBoost™ Poly HRP antibodies (ThermoFisher), followed by tyramide signal amplification (TSA-Dig) using the Opal 6-Plex Detection Kit (Akoya Biosciences). After each round, antibodies were stripped via HIER with Epitope Retrieval Solution 1 (ER1, Leica). Slides were counterstained with DAPI and mounted using ProLong™ Gold antifade reagent (ThermoFisher). Whole-slide images were imported into the HALO image analysis platform (version (4.0), Indica Lab). Cell segmentation was performed using the HALO (HighPlex 4.3.2) module, with nuclei identified based on DAPI staining. Marker positivity thresholds were established using internal positive and negative controls and applied uniformly across all samples. Regions of interest (ROIs) were then manually annotated around immune aggregates and scattered areas and were independently validated by a board-certified hematopathologist blinded to sample identity (controls and MDS samples). MPO staining was used to exclude myeloid populations from evaluation, which might confound analysis due to their endogenous autofluorescence in some channels. Due to significant CD8 staining variability among samples, CD8 staining was excluded from downstream analysis. Quantitative outputs included the total number and frequency of CD20+ B Cells, CD3+ T cells, CD3+CD45RO+ T cells and CD3+CD45RO+TCF1+ T cells within immune aggregates and scattered areas. Data were exported for downstream statistical analysis^56^.

### Multiplex IF Statistical Analyses and Quantification

All MDS and control samples containing at least one immune aggregate and at least one area without immune aggregates containing singly scattered T cells (“scatter regions”) were included for downstream analysis. Cells exhibiting double CD3 and CD20 positivity were excluded from any downstream analysis. The proportions of CD3+ T cells exhibiting TCF1, CD45RO, and double TCF1+CD45RO positivity were calculated for each immune aggregate and across scattered regions within a patient. The difference in proportions for each T cell subset between immune aggregate and scattered region was calculated for each patient. Linear mixed models for the difference in proportion were fit with a fixed effect for either MDS (vs. control), SF3B1 mutation status, or MDS IB1/IB2 status and a random intercept for patient using a compound symmetry correlation structure using the nlme package in R. We tested if the fixed effects of MDS and control were significantly different from zero using the emmeans package in R. Additional details on statistical tests are provided in the corresponding figure legends.

### Spatial TCR and Mutation/Isoform Cell Calling

TCR^+^ cells were identified by first filtering and determining the combination of TCR targeting probes necessary for high patient specificity. Each targeted TCR was patient-specific as determined from scTCR-seq. To determine the patient-specificity of each probe, a predicted false discovery rate (FDR) was first computed for each TCR targeting probe *p* corresponding to patient *s* across probe count thresholds *n*. For each threshold *n*, the expected number of false positive cells in *s* was estimated by scaling the fraction of T/NK cells satisfying the threshold in patient samples that are not from patient *s* by the total number of T/NK cells in samples from patient *s*. Specifically, if *A*_¬*s*_(*n*) and *N*_¬*s*_ denote the number of T/NK cells with at least *n* probes and the total number of T/NK cells in samples not from patient *s*, respectively, and *N*_*s*_ denotes the total number of T/NK cells in samples from patient *s*, the expected number of false positives is calculated as

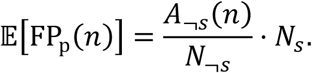

The predicted FDR is then defined as the ratio of the expected number of false positives to the observed number of T/NK cells exceeding the threshold in samples from patient *s*, *A*_*s*_(*n*):

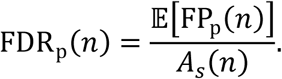

The threshold *n* for each probe *p* was selected as the minimum *n* required to satisfy FDR_p_(*n*) ≤ 0.05. These thresholds were then used to determine TCR^+^ cells annotated as T/NK cells yielding 965 TCR^+^ T/NK cells. Additionally, we also determined that eight paired TCRa and TCRb targets were patient specific when both ≥1 alpha and ≥1 beta probe was detected in a single T/NK cell. For these cases, we also defined TCR^+^ T cells as T/NK cells with ≥1 alpha and ≥1 beta probe. This paired TCRa and TCRb selection yielded 163 T/NK cells that were 100% patient specific. Taking the union of TCR^+^ cells detected with both methods yielded a total of 977 TCR^+^ T/NK cells.

There are cases where non-T/NK cells passed the probe requirement to be called a TCR^+^ cell (**Extended Data Fig. 9b**). Further inspection of these cells revealed the expression of both T cell markers and markers of the assigned cell type (**Extended Data Fig. 9d**), suggesting that these are most likely a product of multiple overlapping cells. Any TCR probes found to associate with non-T cells were excluded; only TCR probes within T/NK cells were considered TCR^+^ cells for downstream analyses.

Similarly, the predicted FDR in mutant samples for each mutant probe at varying probe quantity thresholds was used to determine the ideal threshold for assigning mutant cells. From this, the following probe thresholds were used to annotate mutant cells: ≥1 *ORAI2* alternative isoform, ≥2 SF3B1 mutant, ≥1 SF3B1 mutant + ≥1 *ZDHHC16*, all resulted in predicted FDRs <0.05 for SF3B1 mutant samples; ≥2 U2AF1 mutant resulted in a predicted FDR of 0.021 in U2AF1 mutant samples. This yielded a total of 10,560 SF3B1 mutant cells and 93 U2AF1 mutant cells.

### Alternative splicing analysis from bulk RNA sequencing

Publicly available RNA sequencing from Beat AML (phs001657), TCGA AML (phs000178), Leucegene (GSE232130), and Pellagati et al. (GSE114922) cohorts were used for SF3B1 analyses and compared to normal bone marrow controls from Madan et al. (GSE63816) and Maiga et al. (GSE98310) as well as normal tissue controls from BodyMap2.0 (GSE30611) and the Human Protein Atlas (E-MTAB-1733)^57–63^. For dbGaP data (Beat AML and TCGA AML), BAM files were downloaded from the Genomic Data Commons Portal and converted to FASTQ files using SAMtools (v1.21)^64^. For all samples, reads were trimmed using fastp (v0.23.4) and then aligned to GRCh38 using Spliced Transcripts Alignment to a Reference (STAR, v2.7.11b) in two-pass mode^65,66^. SF3B1 mutations were called from aligned reads using GATK HaplotypeCaller (v4.6.2.0)^67^. Cryptic alternative 3’ splicing events were identified using rMATS (v4.3.0) with novelSS enabled by comparing junction counts of normal controls (n = 63) and SF3B1 mutant patients (n = 76)^68^. Events were filtered to include those with an average change in percent spliced in (ΔΨ) of 10% with a minimum of at least 5 median read counts across mutant samples. For each event, complete coding sequences for both the cryptic and canonical isoforms were constructed by modifying the affected exon boundary within MANE transcript models and removing those that result in frameshift mutations or the insertion of premature termination codons. This was followed by in silico translation using Biostrings (v2.76.0) in R (v4.5.1). Candidate TGFβ-related events were selected by filtering for those in the TGFβ MSigDB Hallmark pathway and validated using the Integrative Genomics Viewer. Sashimi plots were generated using ggsashimi (v1.1.5)^69^.

### TGFBR1 Protein Modeling

Structural modeling of TGFBR1 was performed using PyMOL. The AlphaFold3-predicted wild-type TGFBR1 structure was superimposed onto published crystal structures using residues 24-126 of the extracellular domain (PDB: 9B9F, which also contains TGFBR2 and TGF-β3) and residues 200-503 of the intracellular domain (PDB: 1IAS)^70,71^. The AlphaFold3-predicted mutant TGFBR1 structure containing a four-residue insertion at positions 115-118 was then aligned to the wild-type model using conserved flanking residues.

### *In vitro* Transwell Migration Assay

Isolated bone marrow mononuclear cells (BMMCs) were thawed, washed in PBS w/o Ca or Mg, and resuspended in serum-free RPMI 1640 media (Gibco, catalog #11875093) supplemented with penicillin-streptomycin (Gibco, catalog #15140122), L-glutamine (Gibco, #25030081), MEM non-essential amino acids (Gibco, #11140050), sodium pyruvate (Gibco, #11360070) and β-mercaptoethanol (Gibco, #21985023). The BMMCs (100 μL; 2 × 10^5^ cells/well) were added to 5.0-μm polycarbonate Transwell inserts (Corning, #3421) and allowed to settle for 10 min at 37 °C and 5% CO_2_. 600 μL of serum-free RPMI media containing all of the previously listed supplements and one chemokine at the following concentrations was added to the lower chambers of the Transwell plate: 100 ng/mL CXCL9 (PeproTech, #300-26-20UG), 200 ng/mL CXCL13 (PeproTech, #300-47-20UG), 100 ng/mL CCL19 (PeproTech, #300-29B-20UG), 100 ng/mL CXCL12 (PeproTech, #300-28A-10UG), and 100 ng/mL CCL21 (PeproTech, #300-35A-20UG). The BMMCs were allowed to migrate for 18 h at 37 °C and 5% CO_2_. Migrated cells were harvested from the lower chamber and stained for characterization by spectral flow cytometry with the following anti-human antibodies: CCR7 (BV650, Biolegend, #353233, clone: G043H7), CD34 (Percp, BD Horizon, #340430, clone: 8G12), CD127 (BB700, BD Horizon, #566398, clone: HIL-7R-M21), CD56 (BUV737, BD Horizon, #: 612766, clone: NCAM16.2), CD19 (AlexaFluor488, BD Horizon, #557697, clone: HIB19), CD45 (BV510, BD Horizon, #563204, clone: HI30), CD25 (RY586, BD Horizon, #568124, clone: 2A3), CD11b (BV711, Biolegend, #101241, clone: M1/70), CD11c (BUV605, Biolegend, #301636, clone: 3.9), CD14 (BV570, Biolegend, #301831, clone: M5E2), CD4 (BUV395, BD Horizon, #563552, clone: SK3), CD3 (BUV805, BD Horizon, #612894, clone: SK7), CD45RA (BUV615, BD Optibuild, #751555, clone: HI100), CD8 (BUV563, BD Horizon, #612915, clone: rpa-t8).

### *In vitro* TGFβ Dose Dependence Functional Assay

Primary bone marrow MNCs isolated from untreated MDS patients were thawed, washed in PBS w/o Ca or Mg, and stained with carboxyfluorescein succinimidyl ester (CellTrace, #C34554) for 2 min at room temperature in the dark. The cells were then resuspended in X-VIVO 20 hematopoietic serum-free media (Lonza Biowhittaker, #BW04-448Q) and plated in a 96-well U-bottom plate (Corning, #353072) at a concentration of 2 × 10^5^ cells/well (200 μL/well). The cells were stimulated with CD3/28 human T cell activator Dynabeads (Gibco, #11131D) in a 1:2 bead-to-cell ratio. Recombinant TGF-β1 protein (R&D Systems, #240-B) was added at varied concentrations (0, 0.1, 0.5, 1.0, 5.0, or 10.0 ng/mL) to each well. After incubating for 120 h at 37 °C and 5% CO_2_, the BMMCs were harvested and stained for evaluation of cytokine production by spectral flow cytometry with the following anti-human antibodies: CD3 (BUV805, BD Horizon, #612894, clone: SK7), CD4 (BUV395, BD Horizon, #563552, clone: SK3), CD8 (BUV563, BD Horizon, #612915, clone: rpa-t8), Granzyme B (V450, BD Horizon, #561151, clone: GB11), Perforin (APC-Fire750, Biolegend, #353318, clone: B-D48), IFNγ (PE, Biolegend, #506507, clone: B27), TNFα (RB705, BD Horizon, #570621, clone: Mab11).

### Generation of Cell Lines, Virus Packaging, and Transduction

The K562 human myeloid leukemia cell line was purchased from American Type Culture Collection (ATCC; #CCL-243). All K562 cell lines were maintained in IMDM supplemented with 10% fetal bovine serum. All cell lines were cultured at 37°C and 5% CO2 in the presence of penicillin (100 U/mL) and streptomycin (100 μg/mL). All cell lines were Mycoplasma-free and routinely tested by the Antibody and Bioresource Core Facility at Memorial Sloan Kettering Cancer Center (MSKCC) using the MycoAlert Mycoplasma Detection Kit (Lonza, LT07-701) and MycoAlert Assay Control Set (Lonza, LT07-518).

For generation of TGFBR1 knockout cells, K562 cells stably expressing Cas9 were generated by transduction with an EF1A-Cas9-NLS-HA-P2A-Hygromycin lentiviral vector (Vector Builder) and selected with hygromycin. sgRNAs targeting human TGFBR1 were cloned into a dsRED lentiviral expression vector (Addgene #128055) and tested individually to identify the construct yielding optimal TGFBR1 knockout. TGFBR1 knockout cells were generated by transducing K562-Cas9 cells with dsRED lentiviral vectors encoding sgRNA (TAAAAGGGCGATCTAATGAA; exon 3) targeting human TGFBR1. Transduced cells were selected by fluorescence-activated cell sorting (FACS) for dsRED positivity using a FACSAria III cell sorter (BD Biosciences, RRID:SCR_016695) and TGFBR1 KO cells were confirmed by western blotting. For overexpression and re-expression studies, TGFBR1 isoforms short (NM_004612.4) and long (NM_001306210.2) were cloned into an hPGK-TGFBR1 isoform-P2A-TagBFP2 lentiviral expression vector (Vector Builder) and selected by cell sorting for BFP positivity as described above. Overexpression and re-expression of TGFBR1 isoforms was confirmed by western blotting.

Lentiviral supernatants were produced by transfecting 293T (purchased from ATCC; #CRL-3216) cells with lentiviral constructs and packaging plasmids pVSVG and psPAX2 using PEI. Viral supernatants were collected at 48 and 72 hours post-transfection and used for transduction. Cells were plated at 200,000 cells per well in 24-well plates and resuspended in 1 mL of viral supernatant containing polybrene (4 μg/mL; Sigma-Aldrich, #TR-1003-G). Plates were centrifuged at 2,300 rpm for 90 minutes at 32 °C.

### Western Blots

For western blot analysis, cells were lysed in radioimmunoprecipitation assay (RIPA) buffer (Cell Signaling Technology, #9806S) supplemented with protease (Roche, #11836170001) and phosphatase inhibitor cocktails (Roche, #04906845001). Protein concentration was measured using the BCA Protein Assay Kit (Pierce, #23225). Protein lysates were resolved on 4–12% Bis-Tris gels (Invitrogen, NP0335BOX), transferred to nitrocellulose membranes using the iBlot 2 Gel Transfer Device (Invitrogen, #IB21001), and blocked with Intercept Blocking Buffer (LI-COR, #927-60003). The following primary antibodies were used: TGFBR1 (Abcam, #ab235578), GAPDH (Cell Signaling Technology, #97166), U2AF2 (Cell Signaling Technology, #70471), CD98/SLC3A2 (1:5000; Cell Signaling Technology, #47213), β-actin (1:5000; Cell Signaling Technology, #3700). All primary antibodies were diluted to 1:1000 in Intercept Antibody Diluent (LI-COR) unless otherwise noted. IRDye 800CW donkey anti-rabbit IgG (LI-COR, #926-32213; RRID:AB_621848) and IRDye 680RD donkey anti-mouse IgG (LI-COR, #926-68072; RRID:AB_10953628) secondary antibodies were used at 1:5000 dilution. Bands were visualized using the Odyssey Imager (LI-COR, RRID:SCR_014579).

### Subcellular Fractionation Westerns

Subcellular fractionation was performed using the Subcellular Protein Fractionation Kit for Cultured Cells (Thermo Fisher, #78840) according to the manufacturer’s instructions. Briefly, cells were lysed with the provided fractionation buffers and protein from the cytoplasmic, membrane, and nuclear fractions was isolated. 18 μg of protein per fraction was resolved per well. Transfer, blocking, antibody staining, and imaging were performed as described above.

### Phospho Flow

K562 isogenic cell lines were incubated in serum-free RPMI 1640 for 24 hours prior to cytokine stimulation. Cells were then stimulated with 0 or 10 ng/mL TGF-β1 (Cell Signaling, #75362) for 60 minutes. Following stimulation, cells were stained with anti-CD45 FITC (HI30, #304006, Biolegend) and fixed and permeabilized using Fix/Perm buffer (BD Biosciences) for 10 minutes at 37°C according to the manufacturer’s instructions. Cells were then washed twice with Perm/Wash buffer. Samples were incubated with phospho-SMAD2 (Ser465/Ser467) Alexa Fluor 647 (Cell Signaling Technology, #68550) and analyzed using a BD LSR Fortessa flow cytometer.

### RT-PCR

RNA was isolated from primary MNCs from 5 healthy controls, 5 MDS patients harboring the SF3B1^K700E^ mutation, and 5 MDS patients without splicing factor mutations based on MSK IMPACT testing using the RNeasy Mini Kit with on-column DNase digestion (QIAGEN, #74106). An equivalent amount of RNA from each sample was reverse-transcribed to cDNA using the Verso cDNA Synthesis Kit (Thermo Fisher Scientific, #AB-1453/B). cDNA was then amplified by PCR using the following primer pairs: TGFBR1 Exon 2 Fwd: 5’-CCTCGAGATAGGCCGTTTGT-3’, TGFBR1 Exon 3 Rev: 5’-GCCAGTTCCACAGGACCAA-3’; MAP3K7 Exon 4 Fwd: 5’-TGTCTTGTGATGGAATATGCTG-3’, MAP3K7 Exon 5 Rev: 5’-TCCCTGTGAATTAGCGCTTT-3’; SMURF2 Exon 8 Fwd: 5’-AGCCCTGGCAGACCTCTTAGCT-3’, SMURF2 Exon 9 Rev: 5’-ACCTGGCCTTGTTGCGTTGTCC-3’. PCR was performed under the following cycling conditions: 95°C × 2 min; (95°C × 30 s, 57°C × 30 s, 72°C × 30 s) × 35 cycles; 72°C × 5 min. PCR products were resolved on a 1.5-2% agarose gel and imaged using the Azure 300 Imaging System (Azure Biosystems, RRID:SCR_026671).

### Statistical Analyses

All bar graphs represent the average and error bars represent the standard deviation. All box plots represent the interquartile range (IQR) from the 25^th^ percentile (Q1) to the 75^th^ percentile (Q3) with a line demarcating the median. The lower whisker represents Q1 – 1.5*IQR and the upper whisker represents Q3 + 1.5*IQR. All p-values displayed in bar graphs and boxplots were computed using the Wilcoxon rank-sum test. P-values on paired boxplots were determined with a paired Wilcoxon rank-sum test. The significance symbol values are as follows: ns: p > 0.05, *: p ≤ 0.05, **: p ≤ 0.01, ***: p ≤ 0.001, ****: p ≤ 0.0001. Chi-squared tests were used to assess differences in cell type counts near vs far from mutant cell populations.

## Data and Code Availability

All processed single cell CITE-seq, TCR-seq, and Xenium data will be made publicly available upon peer-reviewed publication. Relevant code will be made publicly available upon peer-reviewed publication.

## ACKNOWLEDGEMENTS

We thank the patients and their families who contributed to this research. We acknowledge DrawImpacts for their contributions to the schematic illustrations. This work was supported by the Neil S. Hirsch Foundation, Edward P. Evans Foundation, the V Foundation, Break Through Cancer, Doris Duke Charitable Foundation Physician Scientist Fellowship, American Society of Hematology Research Training Award for Fellows, American Society of Clinical Oncology Young Investigator Award, the Parker Institute for Cancer Immunotherapy, the MSK Center for Tumor-Immune Systems Biology Pilot, the MSK Gerstner Physician Scholar Program, the National Institutes of Health (1K08CA29327-01, T32 GM132083, R37CA259260, R35CA304457, R01 CA251138, R01 CA283364, R01 CA242020, R01 HL128239, P50 CA254838-01, P30 CA08748), and Blood Cancer United.

## AUTHOR CONTRIBUTIONS

R.F.S., B.D.Z., K.A., O.A.-W., and S.D.W. developed the study concept. C.S., N.F., and R.C. performed spatial molecular analysis using a panel including custom probes designed by R.S., B.D.Z., K.A., M.Z. and S.D.W. R.F.S., B.D.Z., S.D.W, S.M. and K.A. analyzed and interpreted the spatial molecular imaging data. B.D.Z., S.M, and B.D.G. performed computational analysis and developed computational methods for single cell spatial, CITE, and TCR-sequencing data. B.G., K.W., A.M.L., Y.E. designed and performed the single cell CITE and TCR-sequencing. R.F.S., B.D.Z., K.A. analyzed and interpreted the single cell CITE and TCR-sequencing data. K.A., E.R., U.B., N.F., M.R., and A.D. performed histological analyses and guided sectioning and staining. R.F.S., Z.K., K.W., J.W., G.S., R.T., E.M.S., and S.D.W. provided clinical insights and access to patient specimens. R.F.S., K.A., N.A., M.L., L.B., and S.D.W. performed mIF imaging analysis and data interpretation. S.S. performed statistical analyses of mIF. M.M. and J.B. performed analyses of mis-spliced mRNAs. C.P, B.G, and C.W. performed *in vitro* T cells assays under supervision of S.D.W. R.F.S. and P.Z. performed *in vitro* analyses of TGFBR1. R.F.S., B.D.Z., and K.A. generated the tables and figures with guidance from O.A.-W. and S.D.W. Funding for the work was provided by M.v.d.B., B.D.G., O.A.-W., and S.D.W. The study was supervised by B.D.G., O.A.-W., and S.D.W. The manuscript was written R.F.S., B.D.Z., K.A., S.M., O.A.-W., and S.D.W. All authors reviewed the manuscript.

## COMPETING INTERESTS

B.D.G. has received honoraria for speaking engagements from Merck, Bristol Meyers Squibb and Chugai Pharmaceuticals; has received research funding from Bristol Meyers Squibb, Merck and ROME Therapeutics; and has been a compensated consultant for Darwin Health, Merck, PMV Pharma, Shennon Biotechnologies, Synteny and Rome Therapeutics of which he is a co-founder. M.v.d.B. received honorarium from Seres and Garuda; his spouse receives honorarium from Novartis, Synthekine, Beigene, Kite, MustangBio, and Cellectar. He receives royalties from Wolters Kluwer and stock options from Seres and Thymofox. He has IP licensing related to Seres and BMT. He is on the board of DKMS and Smart Immune and external advisory board for the Fred Hutchinson and Fox Chase. O.A.-W is a founder and scientific advisor of Codify Therapeutics, holds equity in this company, and receives research funding from this company. O.A.-W. has served as a consultant for Foundation Medicine Inc., Merck, Prelude Therapeutics, Amphista Therapeutics, MagnetBio, and Janssen, and is on the Scientific Advisory Board of Envisagenics Inc., Harmonic Discovery Inc., and Pfizer Boulder; O.A.-W. has received prior research funding from H3B Biomedicine, Nurix Therapeutics, Minovia Therapeutics, and LOXO Oncology unrelated to the current manuscript. E.S. has received research funding from Eisai Pharmaceuticals and consulting fees for topics unrelated to this manuscript from Abbvie, Agios, Astellas, AstraZeneca, Celgene, Daiichi-Sannkyo, Genentech, Gilead, Jazz, Kura, Servier, and Syndax. R.C. is on the Scientific Advisory Board of Sanavia Oncology and a compensated consultant for the Gerson Lehrman Group. S.D.W. has received research funding from Tigen Pharma and honorarium from Aptitude Health. The remaining authors declare no competing interests.

---

**Extended Data Fig. 1:**
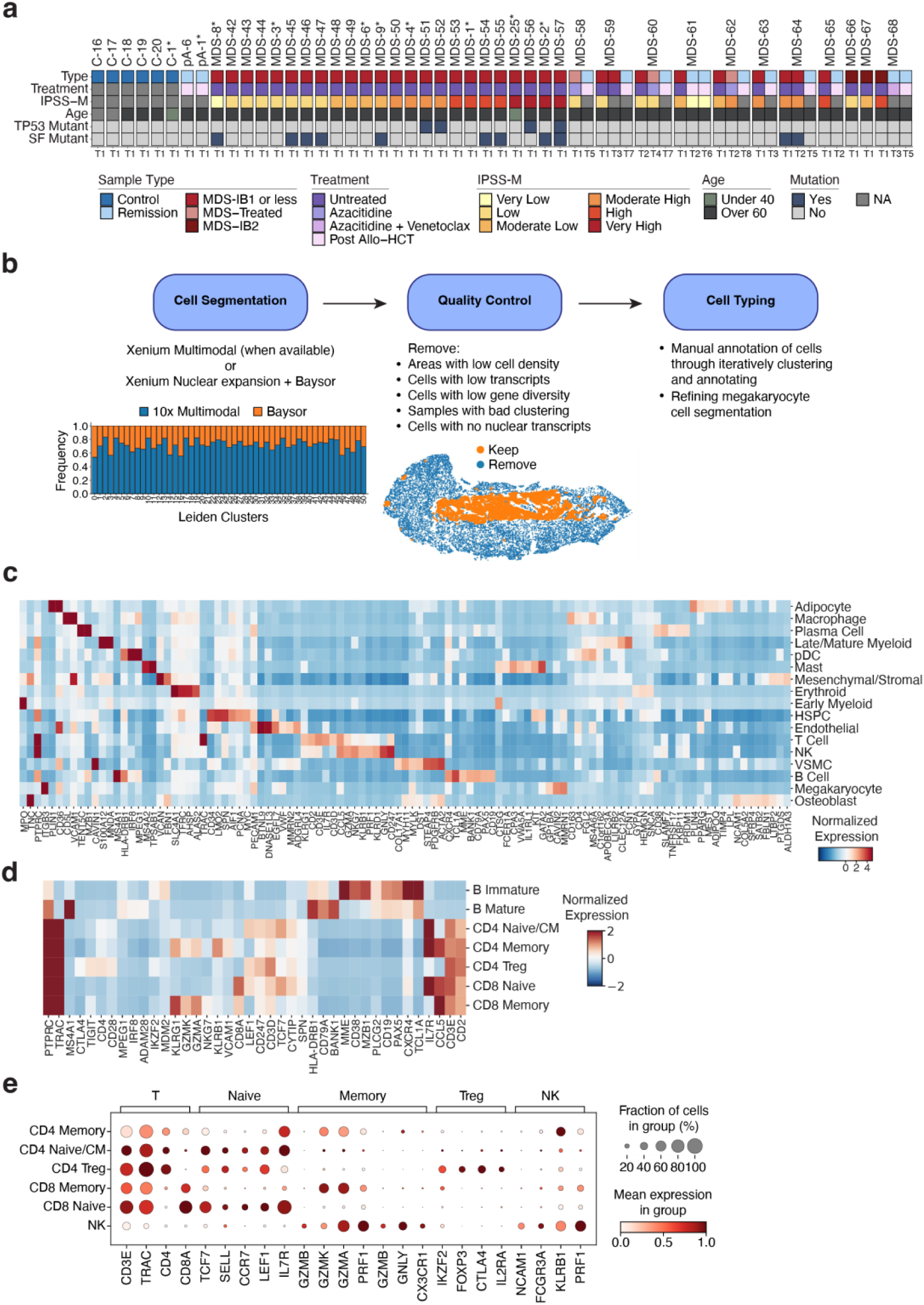
Workflow of cell segmentation and cell typing analysis of human bone marrow specimens. **a**, Oncoprint of patient bone marrow aspirate samples for CITE- and TCR-seq. From top to bottom: Disease status and diagnosis (“Type”), status according to treatment type (“Treatment”), IPSS-M risk category, age, and presence of TP53 (“TP53 Mutant”) or splicing factor (“SF Mutant”) mutation. Asterisks indicate samples whose bone marrow was used for both spatial analysis as well as CITE- and TCR-seq and correspond to spatial cohort sample labeling. **b,** Schematic of workflow of Xenium bone marrow data processing. Initial cell segmentation was performed using the available Xenium multimodal cell segmentation when available (10x Genomics) or Xenium nuclear expansion followed by Baysor re-segmentation. Stacked bar plot displaying proportion of cells attributed to each segmentation method (Multimodal or nuclear expansion + Baysor) by Leiden cluster is included (Leiden clusters 0 to 50 are displayed). Individual samples were then filtered to meet quality control standards, and sparse sample areas were excluded. Refinement of megakaryocyte cell segmentation (see **Methods**) was performed following manual cell typing of marrow populations. **c,** Heatmap showing normalized gene expression scaled by column (gene) of top 10 expressed genes for each cell population in spatial data. **d,** Heatmap showing normalized gene expression scaled by column (gene) of top 15 expressed genes for each adaptive immune cell sub-population. **e,** Dot plots of gene expression markers characteristic of T and NK cell subsets in spatial data.

**Extended Data Fig. 2:**
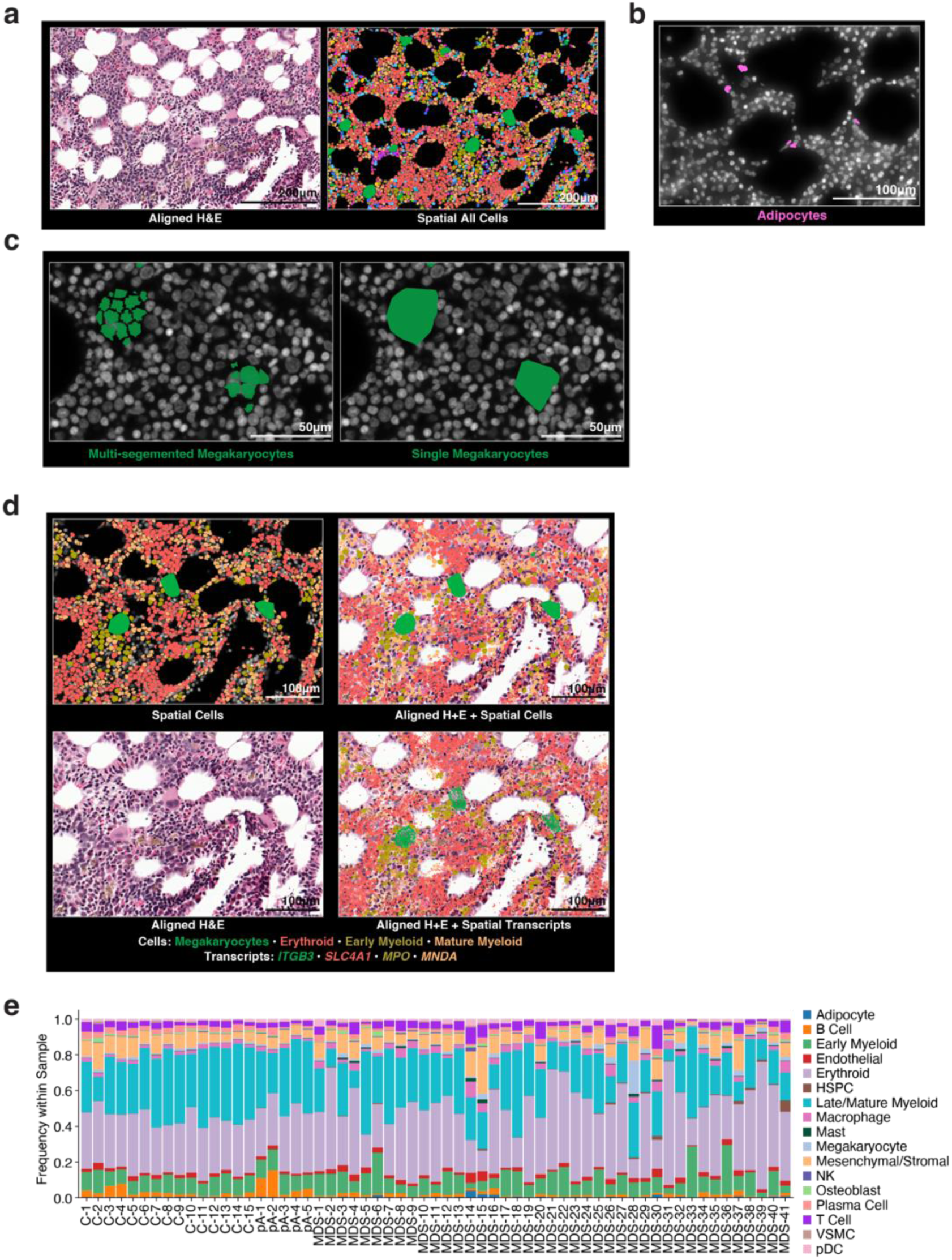
Spatial alignment and refined megakaryocyte segmentation enable accurate cell-type calling and frequency quantification. **a**, Representative images of aligned hematoxylin and eosin (H&E) and Xenium imaging of human bone marrow with cell populations as in **Fig. 1d. b,** Representative image of human bone marrow adipocytes captured by spatial profiling. **c**, Example of specialized megakaryocyte cell segmentation refinement (see **Methods**) for identification of individual megakaryocytes and individual megakaryocyte cell borders. Left: Representative image of initial megakaryocyte segmentation. Right: representative image of refined megakaryocyte segmentation. **d**, Example of spatial cell populations and transcripts projected onto post-Xenium H&E of the same area. Top left: spatial image of cell populations by color based on cell segmentation as depicted. Bottom left: aligned H&E. Top right: projection of Xenium cell populations over aligned H&E. Bottom right: projection of cell type specific transcripts onto aligned H&E. **e,** Stacked bar plots of individual cell type frequencies across all bone marrow spatial specimens stratified by patient.

**Extended Data Fig. 3:**
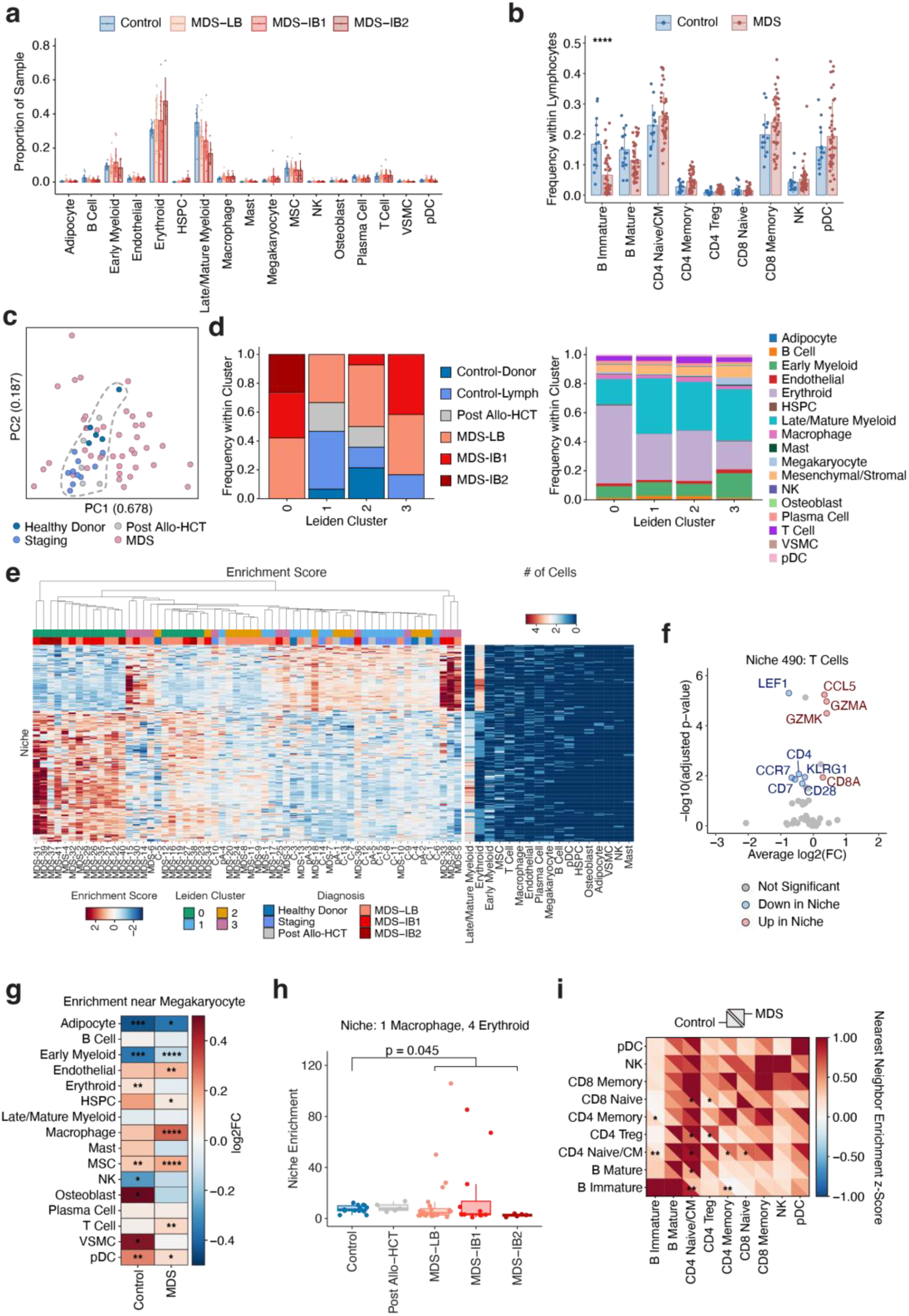
Spatial cell type composition and organization distinguish control from MDS bone marrow. **a**, Bar plot showing average spatial cell type frequencies across controls and MDS disease stages from spatial data. **b**, Bar plot showing average spatial frequencies of lymphocyte subsets in controls and MDS from spatial data. **c**, Principal component analysis visualization of spatial samples computed on sample cell type frequencies colored by sample type. The dotted line encircles all non-MDS samples. **d**, Composition of each Leiden cluster computed from niche enrichment scores colored by sample type (left) and cell type composition of samples (right). **e**, Left, heatmap displaying niche enrichment scores of differentially enriched niches of Leiden cluster 0. Each column is an individual sample, and each row is a niche, each ordered with hierarchical clustering. The top color bars annotate the Leiden cluster and sample type of each sample. Right, heatmap displaying the cell type composition of niches corresponding to each row of the right heatmap. **f,** Volcano plot of differential gene expression between T cells located within niche #490 (niche containing 1 T Cell, 1 MSC, 1 Mk, and 2 Myeloid cells) and T cells that do not reside in niche #490 in MDS samples. **g**, Heatmap displaying spatial enrichment near megakaryocytes for each cell type. Enrichment is calculated by computing the average log2(fold change) between the frequency of each cell type out of all cells that are near a megakaryocyte (≤15µm from a vertex of a megakaryocyte cell polygon) vs far over control (left column) and MDS (right column) samples. Asterisks denote p-values computed from paired Wilcoxon signed-rank test: *p ≤ 0.05, **p ≤ 0.01, ***p ≤ 0.001, ****p ≤ 0.0001. **h**, Box plot of niche enrichment scores of a differentially enriched niche (Leiden cluster 0 samples vs rest) consisting of one macrophage and four erythroid cells. The p-value comparing the enrichment scores of this niche between control and MDS samples is calculated with a Wilcoxon rank-sum test. **i**, Neighborhood enrichment z-score heatmap between pairs of T, B, NK, and pDC cell sub-populations (squares) from control (bottom triangle) and MDS (top triangle) patients. Red indicates positive enrichment (positively associated, >0), blue indicates negative enrichment (negatively associated, <0). Asterisks denote p-values computed from Wilcoxon rank-sum test: *p ≤ 0.05, **p ≤ 0.01.

**Extended Data Fig. 4:**
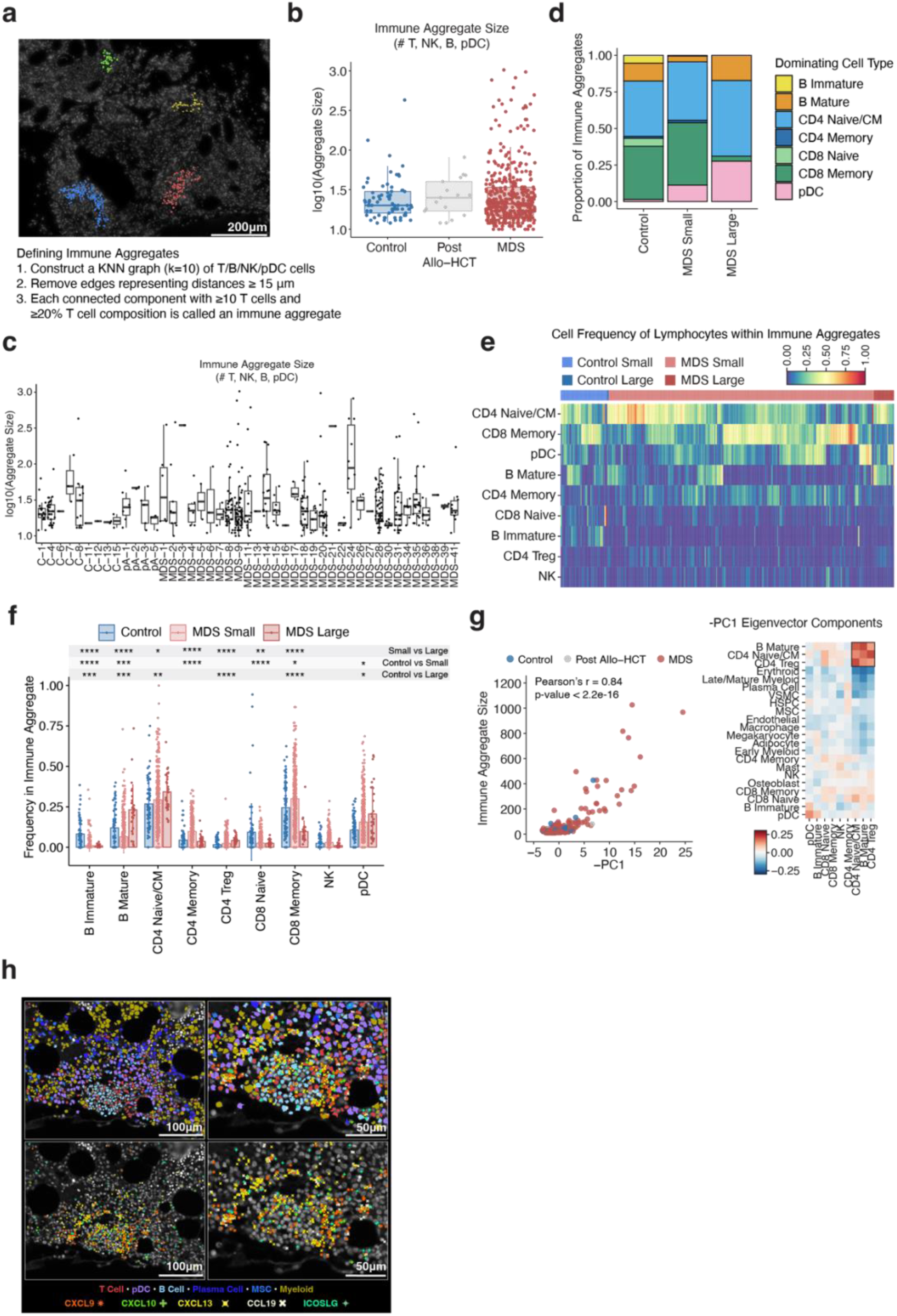
Immune aggregate composition and organization in MDS bone marrow. **a**, Computational definition of immune aggregates and representative image of individual immune aggregates within a human bone marrow sample. Each colored aggregate represents one immune aggregate. **b**, Box plot of immune aggregate sizes grouped by sample type. **c**, Box plots of immune aggregate sizes grouped by individual patient specimen. Box represents interquartile range, line represents median, lower whisker represents Q1-1.5*IQR and upper whisker represents Q3 + 1.5*IQR. **d**, Stacked bar graph of the frequency of immune aggregates as defined by the dominant lymphoid cell type within each immune aggregate across control, MDS small (<100 cells), and MDS large (≥100 cells) immune aggregates. **e**, Heatmap of the cell frequency of lymphocytes within immune aggregates in control, MDS small (<100 cells), and MDS large (≥100 cells) immune aggregates. **f**, Average cell frequency of all lymphoid subsets within control, MDS small (<100 cells), and MDS large (≥100 cells) immune aggregates. Bars represent the average, error bars represent standard deviation. Asterisks denote p-values computed from Wilcoxon rank-sum test: *p ≤ 0.05, **p≤ 0.01, ***p≤ 0.001, ****p≤ 0.0001. **g**, Left, scatter plot: X-axis, -PC1 from PCA of individual immune aggregates using pairwise cell type neighborhood enrichment scores (of T, B, NK, and pDC cells). Y-axis, size of immune aggregates based on quantity of lymphocytes in control and MDS aggregates. Right, heatmap of -PC1 eigenvector components. **h**, Representative images of tertiary lymphoid structure-associated cytokines in and around an MDS immune aggregate. Top left: immune aggregate with each cell type represented by a different color. Bottom left: identical image showing individual cytokine transcripts represented by symbols and colors. Top right: higher magnification of same immune aggregate with cell types (colors) and individual cytokine transcripts (colored symbols). Bottom right: identical image showing individual cytokine transcripts represented by colored symbols.

**Extended Data Fig. 5:**
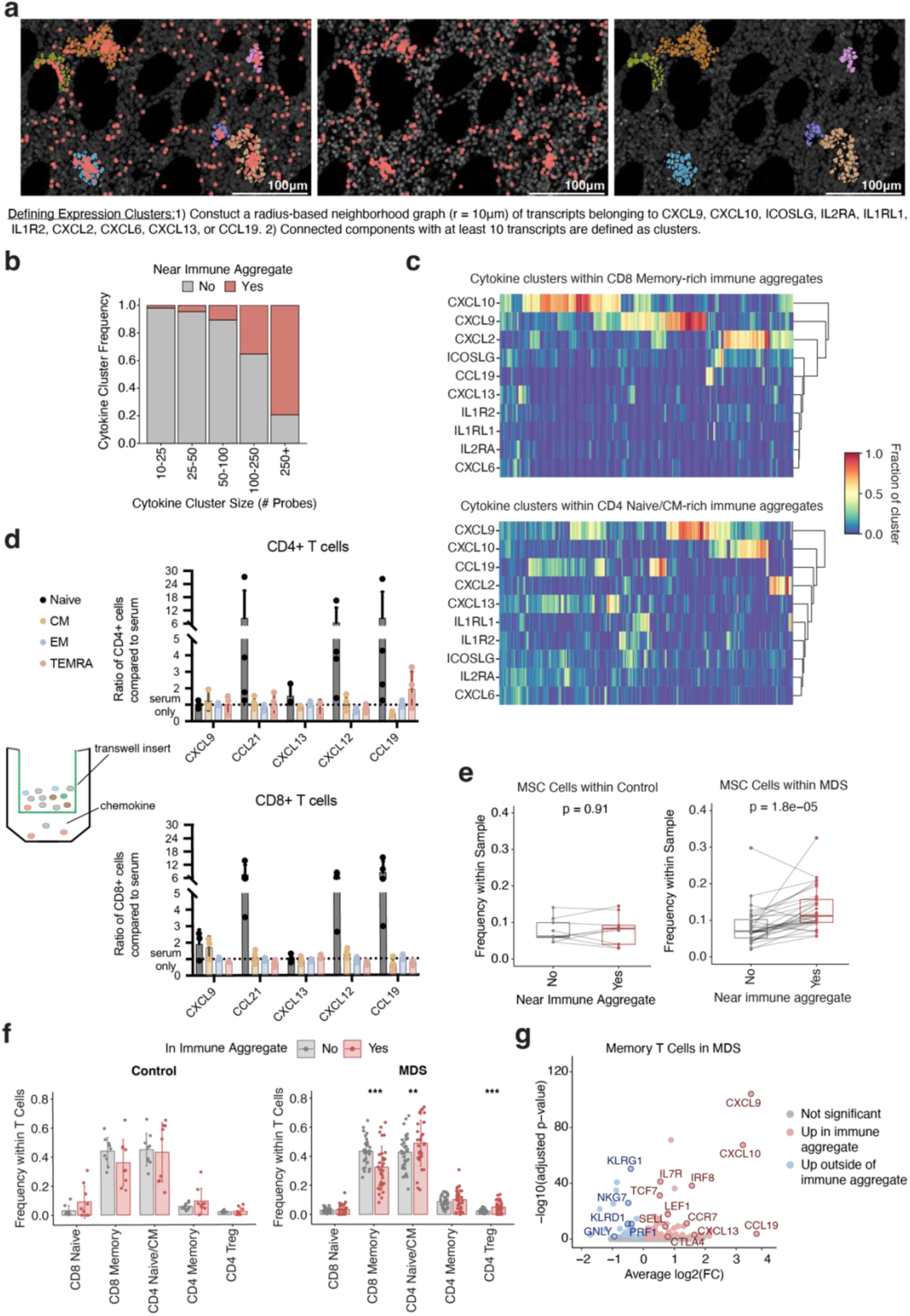
Spatially resolved cytokine and chemokine clusters define immune aggregate microenvironments and T cell migration in MDS. **a**, Computational definition of cytokine and chemokine clusters and representative images of cytokine/chemokine gene clusters (red dots) with associated cell clusters (various colors) within a human bone marrow sample. **b**, Bar plots displaying frequency of cytokine clusters colocalized with cells within immune aggregates stratified by cytokine cluster size as defined by number of cytokine gene probes in the cluster. **c**, Heatmap displaying the composition of cytokine clusters (columns) colocalized with CD8 Memory-rich (≥33% CD8 Memory, top) and CD4 Naïve/CM-rich (≥33% Naïve/CM, bottom) immune aggregates (columns). **d**, Bar plots of transwell assay results showing the mean ratio of migrated CD4+ and CD8+ T cells from n=4 MDS patients following incubation with indicated chemokines, normalized to serum-only control compared to serum only. Mean ± SD. **e**, Paired box plots comparing frequency of mesenchymal/stromal cells (MSCs) within cells near (≤15µm from a T, B, NK, or pDC within an immune aggregate) and far from immune aggregates in control (left) and MDS (right) marrows. P-values were calculated by paired Wilcoxon rank-sum test. **f**, Average relative frequency of T cell subsets outside (No) versus inside (Yes) control and MDS immune aggregates for each patient sample. P-value calculated by Wilcoxon rank-sum test. **: p ≤ 0.01, ***: p ≤ 0.001. **g,** Volcano plot of differential gene expression between memory T cells within MDS immune aggregates (red) versus outside immune aggregates (blue) in MDS samples.

**Extended Data Fig. 6:**
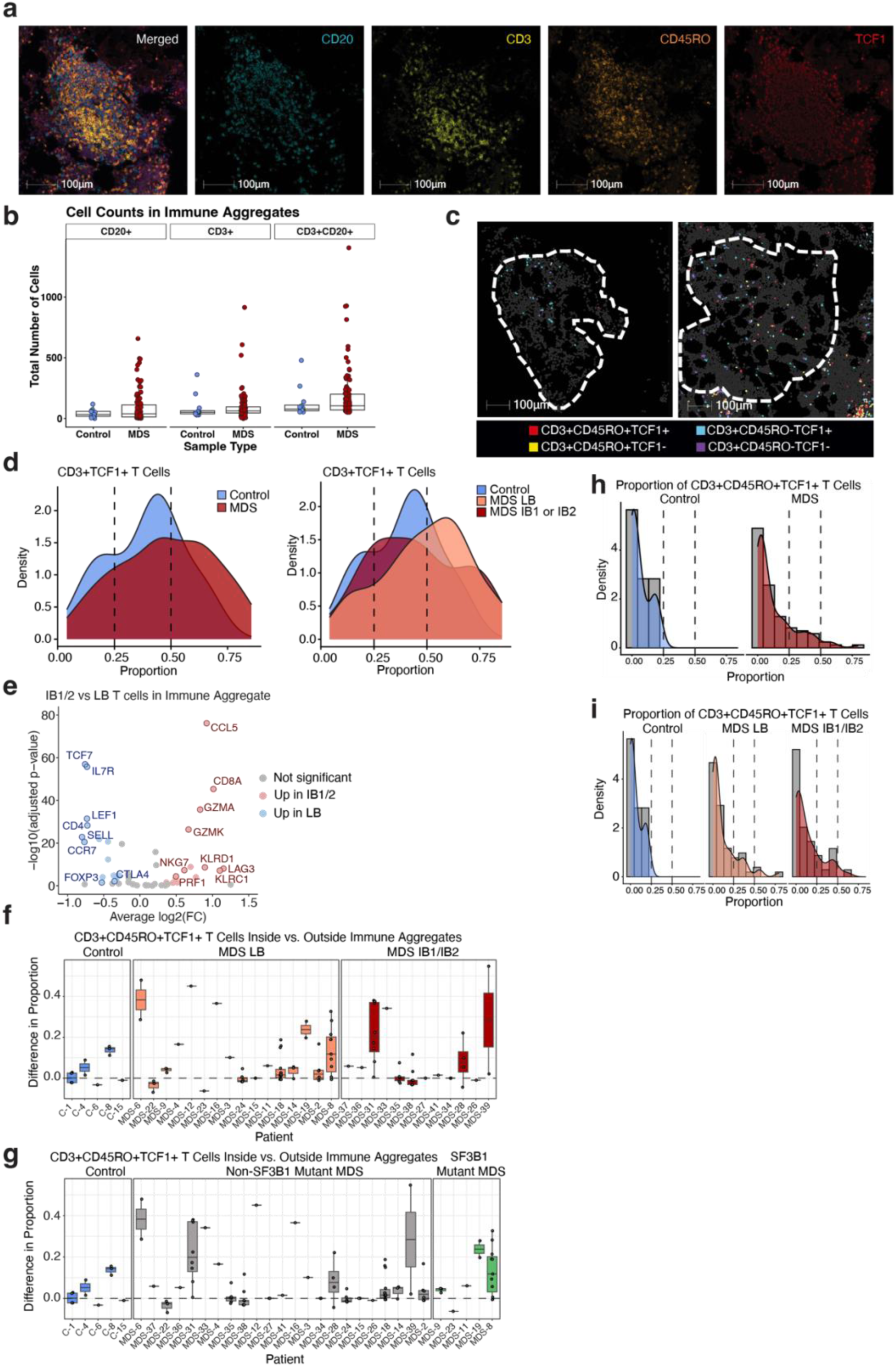
Multiplex immunofluorescence analysis of T cells within MDS immune aggregates. **a**, Multiplex immunofluorescence photomicrograph of an MDS immune aggregate; from left to right: merged image, and single channel images for CD20 (cyan), CD3 (yellow), CD45RO (orange), TCF1 (red). **b,** Total counts of CD3+ T cells and CD20+ B cells across immune aggregates observed in healthy control and MDS samples. Each colored dot represents an immune aggregate. Statistical comparison of the difference in the distributions of these counts between groups shows no statistically significant difference (Wilcoxon rank sum test, p>0.05). **c**, Representative images containing masks of different T cell immunophenotypes in bone marrow areas containing single scattered T cells in a control marrow (left) and MDS marrow (right); CD3^+^CD45RO^+^TCF1^+^ (red), CD3^+^CD45RO^+^TCF1^-^ (yellow), CD3^+^CD45RO^-^TCF1^+^ (cyan), CD3^+^CD45RO^-^TCF1^-^ (purple). **d**, Histogram plots depicting the estimated density of the proportion of each T cell subset across immune aggregates in control and MDS marrows. The denominator is the total number of T cells. Vertical lines denote frequencies of CD3^+^TCF1^+^ subsets at 25% and 50% frequency, respectively. Statistical comparison of the difference in the distributions of these counts within immune aggregates between groups shows no statistically significant difference (Wilcoxon rank sum test, p>0.05). **e**, Volcano plot of differentially expressed genes in T cells within immune aggregates of MDS-LB vs MDS-IB1/2 (data from spatial transcriptomic analysis). **f**, Box plots showing the difference in the proportion of CD3^+^CD45RO^+^TCF1^+^ T cells in immune aggregates versus scatter regions in controls, MDS-LB, and MDS-IB1/IB2 core biopsies (MDS-LB: p=0.001; MDS-IB1/IB2: p=0.028). **g**, Box plots showing the difference in the proportion of CD3^+^CD45RO^+^TCF1^+^ T cells in immune aggregates versus scatter regions in controls, SF3B1-mutated and non-SF3B1 mutant MDS core biopsies (SF3B1 mutated: p=0.128; SF3B1 unmutated; p=0.001). **h** and **i**, Distribution of the proportion of CD3^+^CD45RO^+^TCF1^+^ subsets out of total T cells across all immune aggregates. Vertical lines denote frequencies of CD3^+^CD45RO^+^TCF1^+^ subsets at 25% and 50%, respectively. Statistical comparison of the difference in the distributions of these counts within immune aggregates between groups shows no statistically significant difference when comparing MDS versus healthy control marrows (Wilcoxon rank sum test, p>0.05) or MDS-LB and MDS-IB1/2 (Kruskal-Wallis rank sum test, p>0.05).

**Extended Data Fig. 7:**
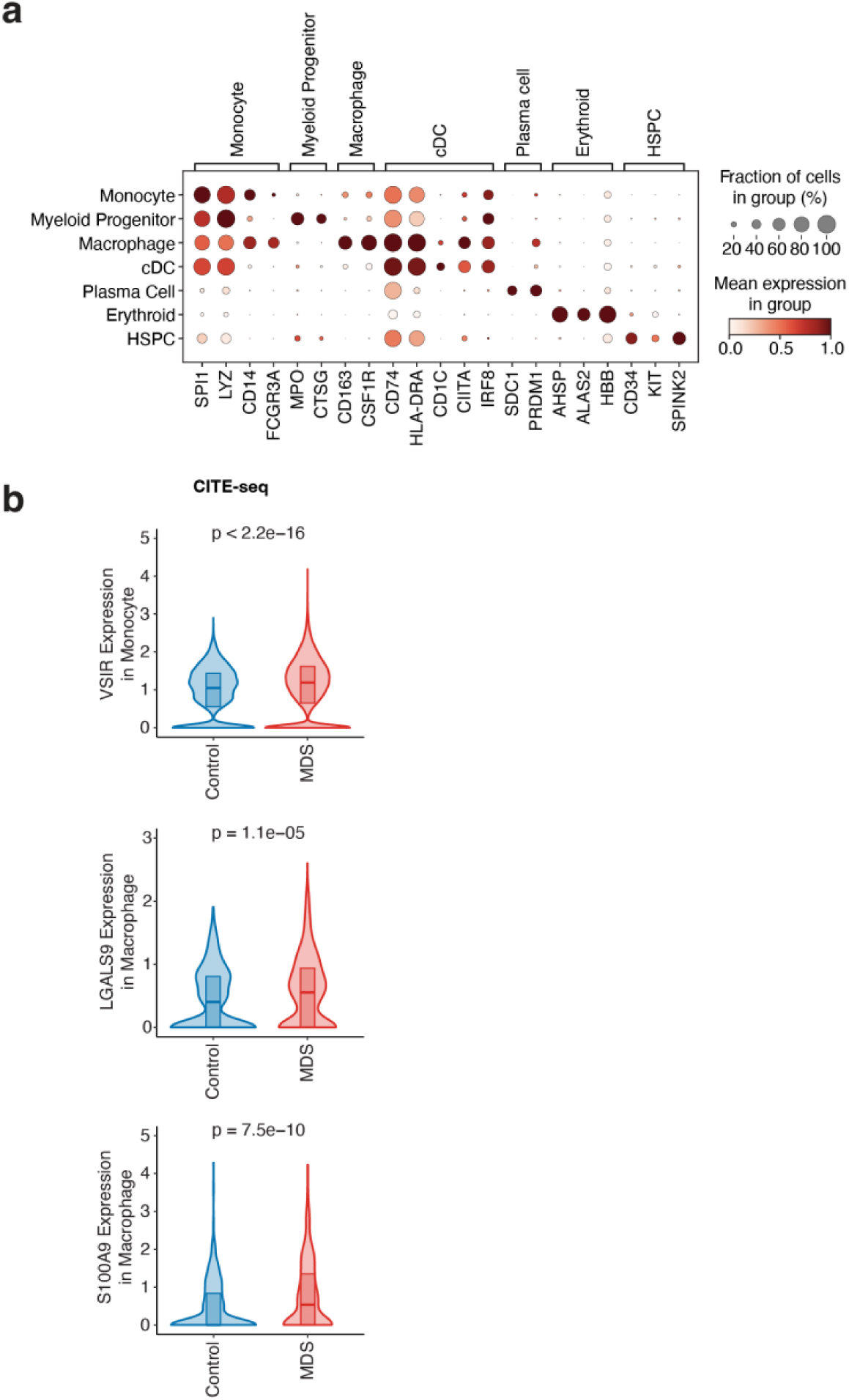
Expression of characteristic markers of myeloid subsets and myeloid immunosuppressive genes in CITE-seq data. **a,** Dot plots of mRNA expression of markers characteristic of CD45dim bone marrow cell populations in healthy control and MDS patient bone marrow cells from CITE-seq data. **b**, Violin plots of myeloid immunosuppressive genes in control and MDS patient bone marrow cells from CITE-seq data. P-values calculated from Wilcoxon rank-sum test comparing to control samples (blue) and MDS samples (red). HSPC, hematopoietic stem and progenitor cell; cDC, conventional dendritic cell.

**Extended Data Fig. 8:**
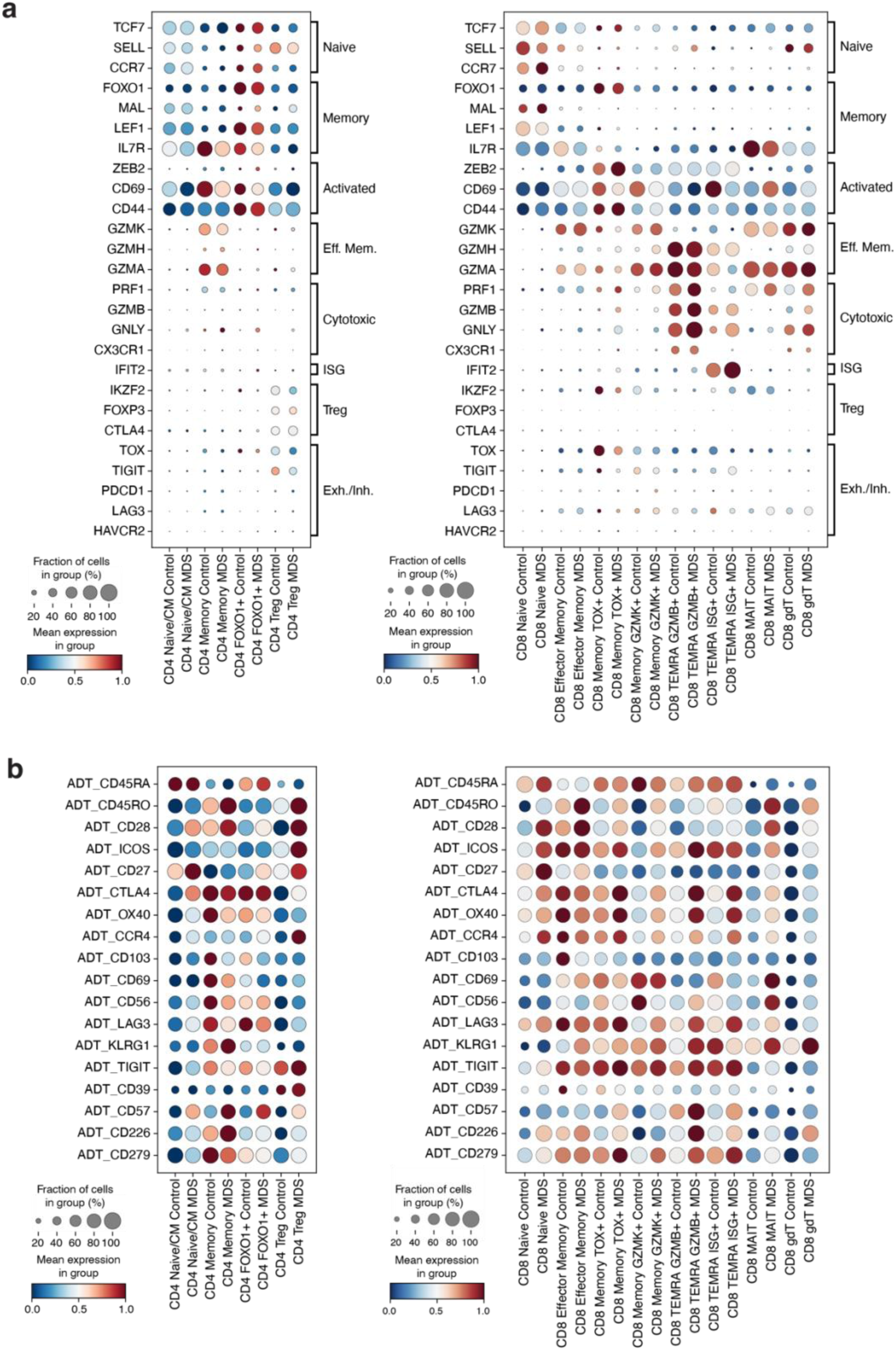
Expression of characteristic markers of CD4 and CD8 T cells at mRNA and protein level from CITE-seq data. Dot plots of **(a)** mRNA expression and **(b)** ADT expression of markers characteristic of CD4 (left) and CD8 (right) T cell subsets in healthy control and MDS patient bone marrow cells from CITE-seq data.

**Extended Data Fig. 9:**
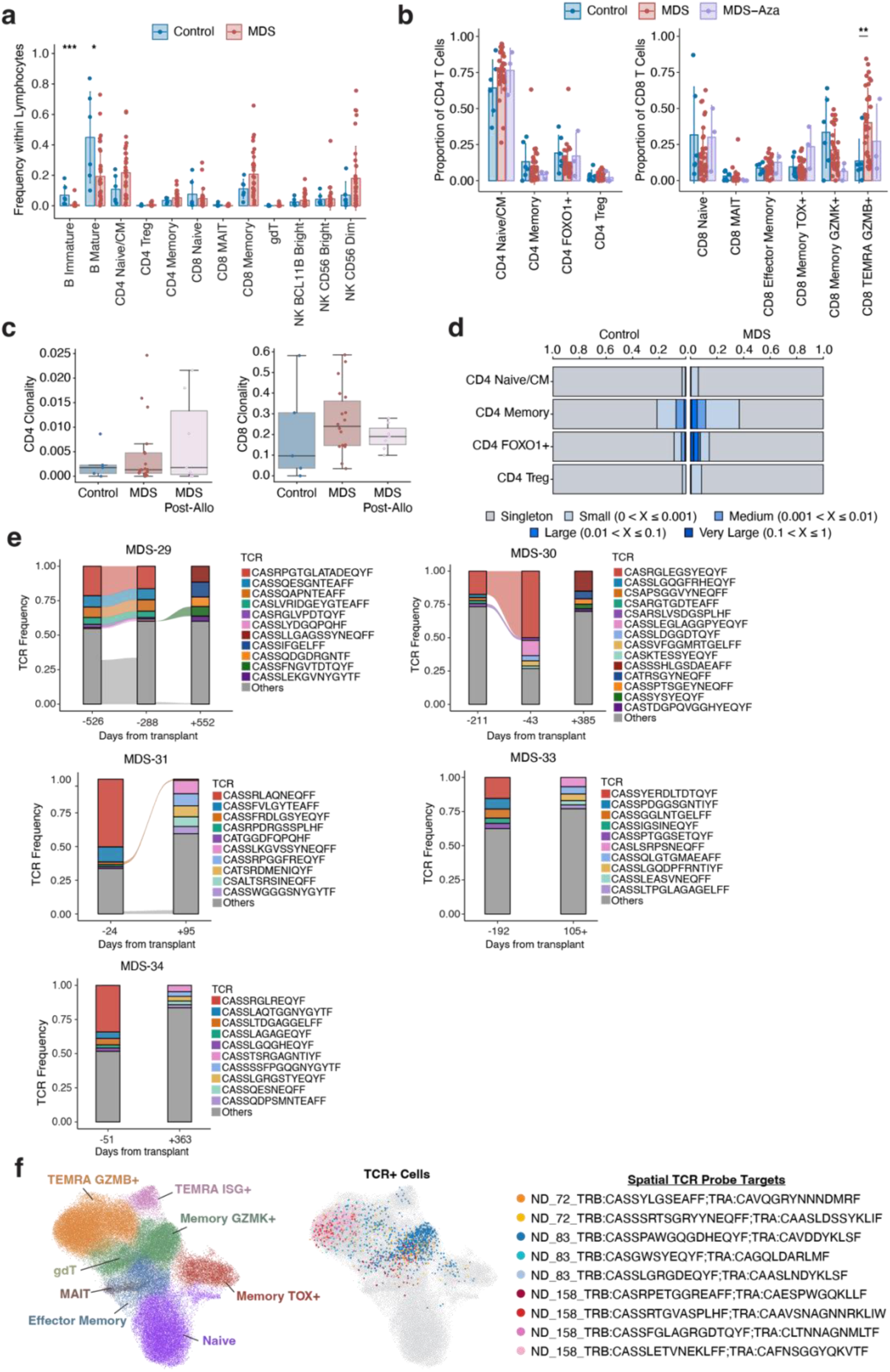
Evaluation of T cell receptors (TCRs) within CITE/TCR-seq of bone marrow aspirates from MDS patients. **a,** Frequency of lymphocytes in healthy control (n=6) and MDS (n=34) patient marrow aspirates. Bars represent average, error bars represent standard deviation. Asterisks denote p-values calculated from Wilcoxon rank-sum test: * p ≤ 0.05, *** p ≤ 0.001. **b**, Frequency of CD4 (left) and CD8 (right) T cell subsets across control (n=6), MDS (n=34), and azacitidine-exposed MDS patient samples (n=3). Asterisks denote p-values calculated from Wilcoxon rank-sum test: **: p ≤ 0.01. **c**, Box plots of CD4 (left) and CD8 (right) T cell clonality in samples with at least 50 CD4 T cells with a defined TCR (n=5 controls, n=19 MDS, n=7 post allogenic transplant) and at least 50 CD8 T cells with a defined TCR (n=5 controls, n=18 MDS, n=7 post allogenic transplant). Box represents interquartile range, line represents median, lower whisker represents Q1-1.5*IQR and the upper whisker represents Q3 + 1.5*IQR. **d**, Clonal frequency of TCRs in CD4 T cell subsets from healthy control (left) and MDS (right) patient bone marrow samples represented by a range of clone sizes from singletons to very large. **e**, Alluvial plots showing TCR frequency of five MDS patients across different timepoints. Largest clones are colored. Negative and positive values represent days from transplant whereby the infusion date is day +1. **f,** Left, UMAP representation of CD8 T cell subsets from CITE-seq data. Middle, UMAP representation of CD8 T cells colored by the top 3 TCRs from 3 MDS patients targeted for spatial detection. Right, list of TCRs (n=3/patient) detected spatially across 3 MDS patients.

**Extended Data Fig. 10:**
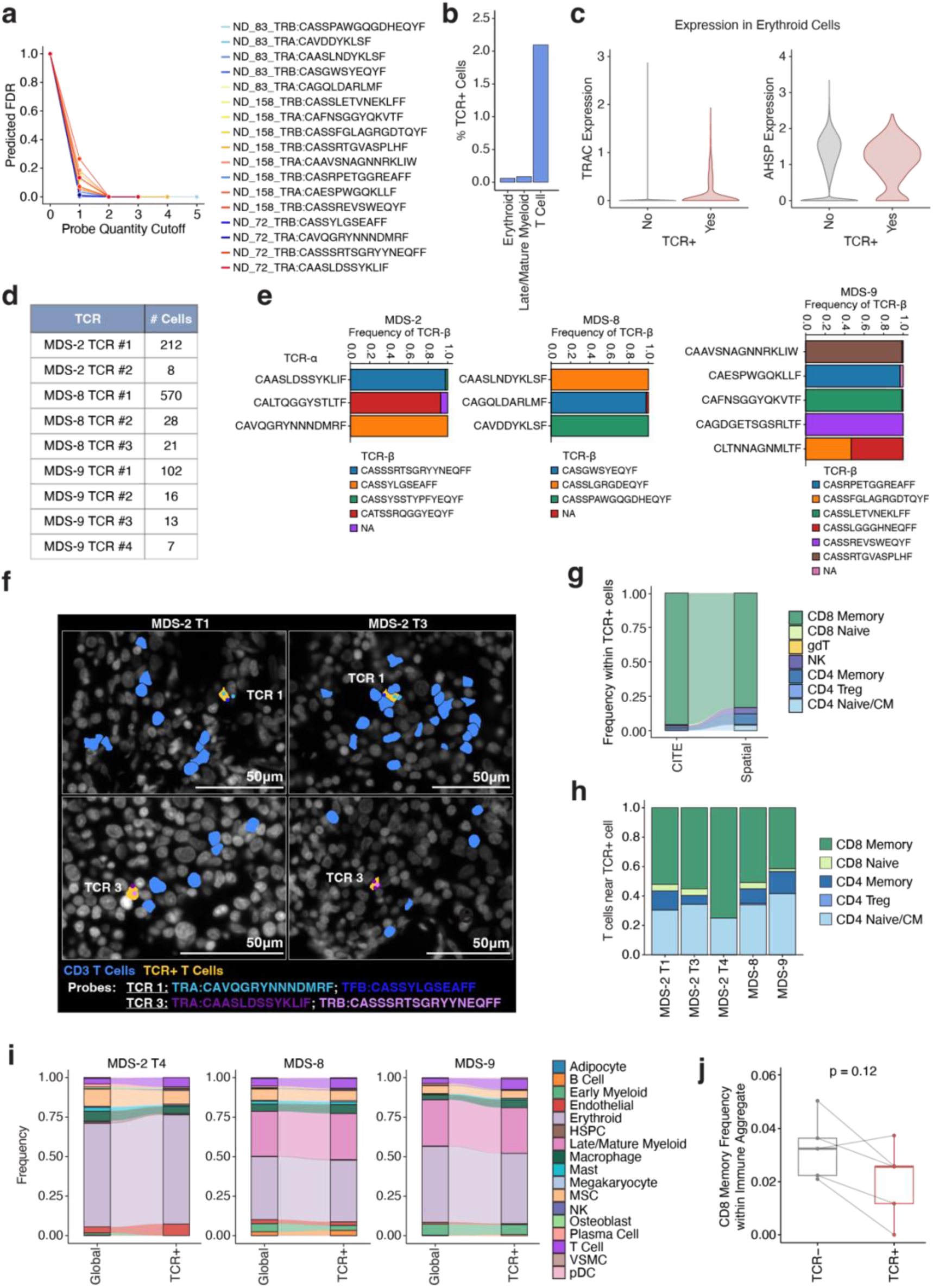
Single cell spatial detection of MDS TCR clones in situ. **a,** Predicted false discovery rate (FDR) of spatial TCR detection based on number of Xenium probes in cell. This is used to determine the probe quantity requirement for a cell to be called “TCR^+^” (see **Methods**). **b**, Bar plot displaying the percentage of TCR^+^ cells within each cell type for the top three cell types with the greatest quantity of TCR^+^ cells. **c**, Violin plots comparing *TRAC* (left) and *ASHP* (right) expression between erythroid cells that pass the TCR probe quantity requirement (“Yes”) and those that do not pass the requirement (“No”). **d,** Table showing the number of TCR^+^ T cells per patient. **e**, Stacked bar plots displaying frequency of TCR-beta sequences associated with each targeted TCR-alpha sequence in scTCR-seq data. **f,** Representative images of T cells (blue) and TCR^+^ T cells (yellow) with TCR probes (colored dots) across T1 and T3 bone marrow core biopsy timepoints from the same MDS patient. **g**, Alluvial plots highlighting the correlation between TCR^+^ T cell phenotypes between CITE-seq and spatial transcriptomic data. **h**, Stacked bar plots displaying frequency of each T cell subtype within 15 µm of a TCR^+^ T cell for each sample. **i**, Alluvial plots comparing global cell type frequencies to cell type frequencies within 15 µm of a TCR^+^ T cell. **j**, Paired box plots comparing frequency of TCR^+^ cells within immune aggregates to CD8 memory T cells within immune aggregates that are not TCR^+^ from n=5 MDS patient samples (MDS-2 T1, T3, T4, MDS-8, and MDS-9). P-value was calculated by paired Wilcoxon rank-sum test. Box represents interquartile range, line represents median, lower whisker represents Q1-1.5*IQR and the upper whisker represents Q3 + 1.5*IQR.

**Extended Data Fig. 11:**
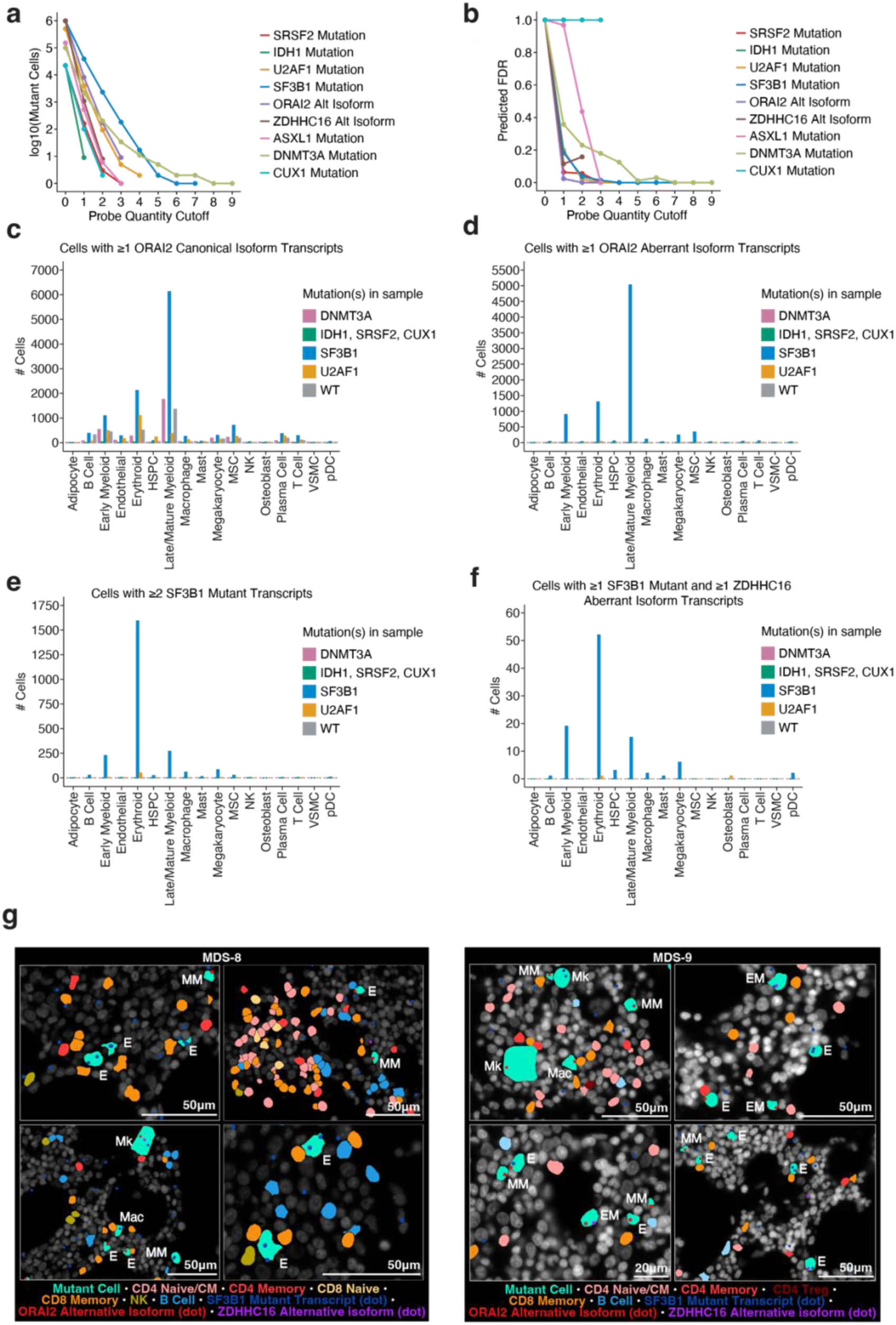
Analysis of somatic mutations in bone marrow core biopsies *in situ* using single cell spatial analysis. **a**, Line graph of the number of mutant cells (Y-axis) according to total probe quantity threshold per cell (X-axis) for each mutation and RNA isoform assayed. **b**, Line graph of predicted false discovery rate (FDR, Y-axis) according to total probe quantity threshold per cell (X-axis) for each mutation and RNA isoform assayed. **c**, Number of cells with ≥1 normal annotated *ORAI2* isoform probe across each cell type grouped by sample mutation status across 16 (n=3 control, n=13 MDS) patient samples. **d**, Number of cells with ≥1 SF3B1-mutant specific *ORAI2* aberrant RNA isoform probe across each cell type grouped by sample mutation status across 16 (n=3 control, n=13 MDS) patient samples. **e**, Number of cells with ≥2 SF3B1 mutant probes across each cell type grouped by sample mutation status across 16 (n=3 control, n=13 MDS) patient samples. **f,** Number of mutant cells identified using the combination of ≥1 SF3B1 mutant probe plus ≥1 aberrant *ZDHHC16* isoform probe across each cell type grouped by sample mutation status across 16 (n=3 control, n=13 MDS) patient samples. **g,** Representative images of SF3B1 mutant cells (turquoise) with detection of ≥2 SF3B1 mutant probes (red dots), ≥1 aberrant *ORAI2* isoform (blue dot) or combination of ≥1 SF3B1 mutant probe (red dot) plus ≥1 aberrant *ZDHHC16* isoform (purple dot) in MDS-8 and MDS-9 patient bone marrows. Mutant cell types are labeled in white. MM, mature myeloid; EM, early myeloid; E, erythroid; Mk, megakaryocyte; Mac, macrophage.

**Extended Data Fig. 12:**
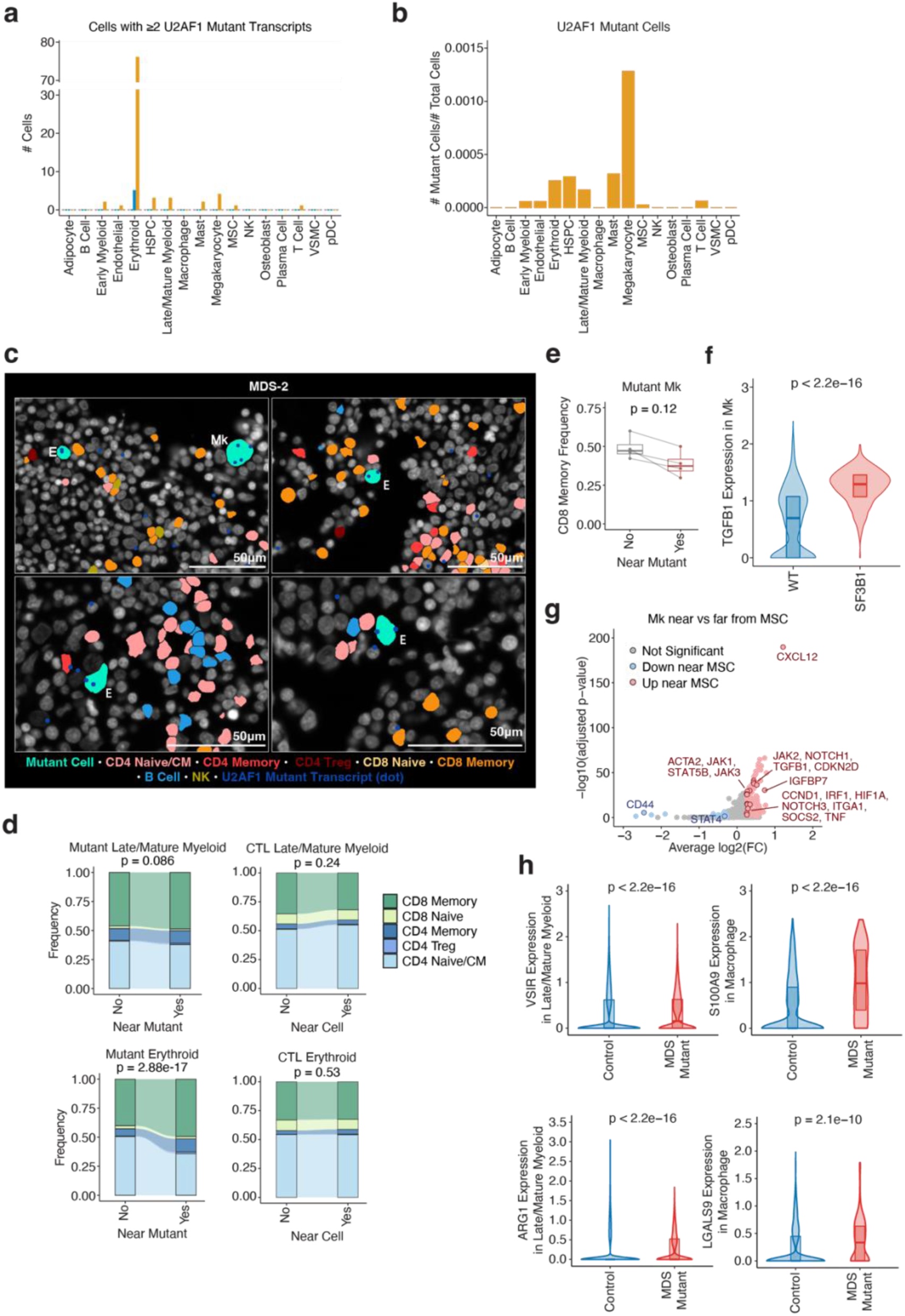
Spatial detection of MDS mutant cells and their surrounding immune microenvironment *in situ*. **a**, Number of cells with ≥2 U2AF1 mutant probes across each cell type grouped by sample mutation status across 16 (n=3 control, n=13 MDS) patient samples. **b**, Proportion of mutant U2AF1 mutant cells in each cell type across n = 5 mutant U2AF1 MDS samples. **c**, Representative images of U2AF1 mutant cells (turquoise) with detection of ≥2 U2AF1 mutant probes (blue dots) in MDS-2 patient bone marrow. Mutant cell types are labeled in white. E, erythroid; Mk, megakaryocyte. **d**, Left, alluvial plot comparing the frequency of T cell subtypes between T cells far (>15 µm) from a mutant myeloid (top) or erythroid (bottom) cell and T cells near (within 15 µm) a mutant myeloid (top) or erythroid (bottom) cell in mutant SF3B1 MDS samples. Right, alluvial plot comparing the frequency of T cell subtypes between T cells far (>15 µm) from a myeloid (top) or erythroid (bottom) and T cells near (within 15 µm) a myeloid (top) or erythroid (bottom) cell in control samples. P-values are calculated with a chi-squared test on T cell subtype counts between “No” and “Yes” groups. **e**, Paired box plots comparing frequency of CD8 Memory cells out of all T cells near (within 15 µm) a mutant Mk and the frequency of CD8 Memory cells out of all T cells far (>15 µm) from a mutant Mk. P-value was calculated by paired Wilcoxon rank-sum test. Box represents interquartile range, line represents median, lower whisker represents Q1-1.5*IQR and upper whisker represents Q3 + 1.5*IQR. **f**, Violin plots comparing *TGFB1* expression in SF3B1 mutant megakaryocytes (“SF3B1”) and megakaryocytes from control samples (“WT”). P-value was calculated using a Wilcoxon rank-sum test. Internal box represents interquartile range, and the line represents the median. **g**, Volcano plot comparing gene expression of megakaryocytes near (within 15um from megakaryocyte polygon vertex to centroid point of MSC) or far (> 15um from MSCs) from an MSC (n=13 samples from 9 patients). **h**, Violin plots comparing gene expression of *VSIR*, *LGALS9*, *ARG1*, and *S100A9* in mutant myeloid cell populations compared to myeloid cells in control samples. P-value was calculated using a Wilcoxon rank-sum test. Internal box represents interquartile range, and the line represents the median.

**Extended Data Fig. 13:**
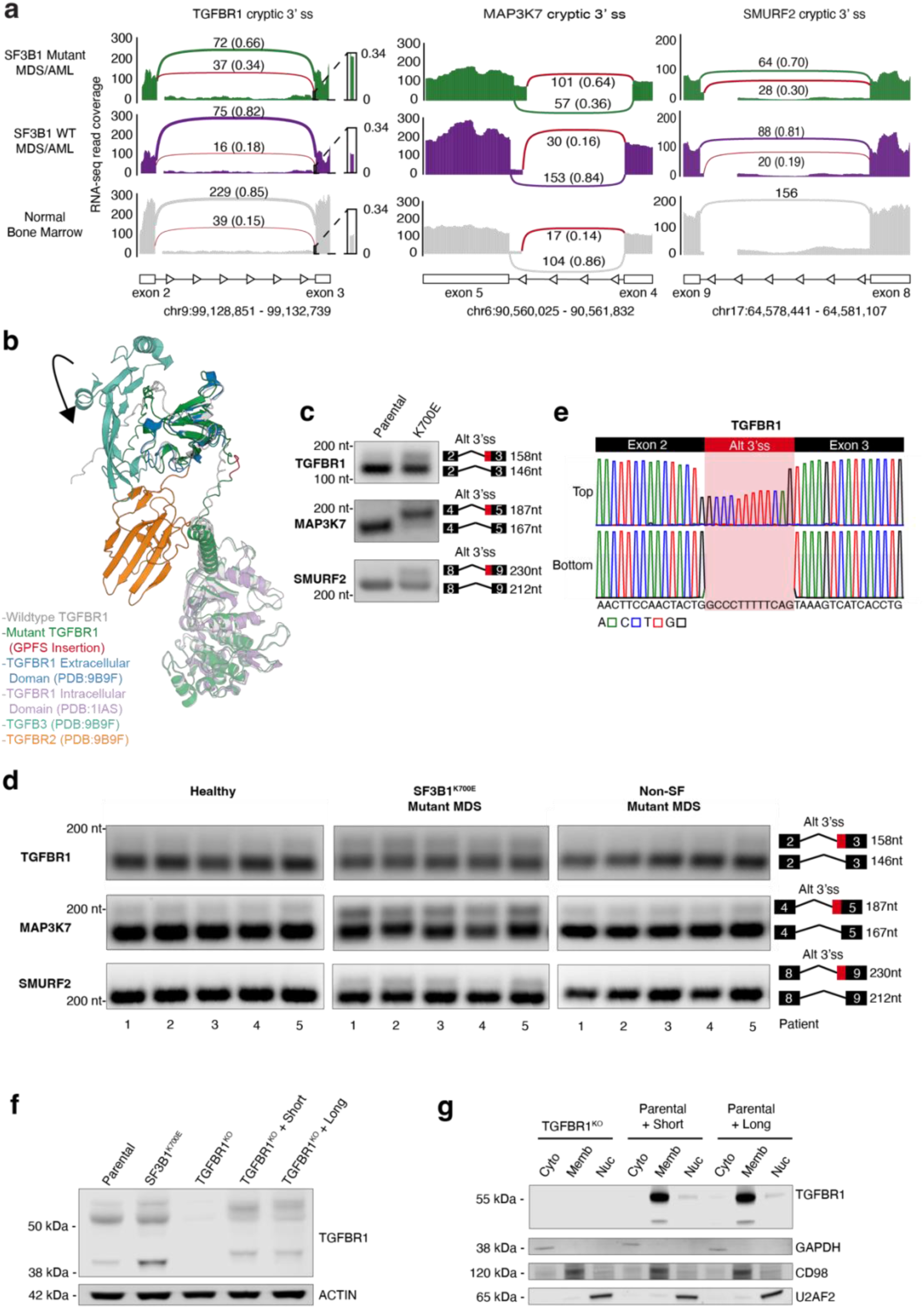
Altered splicing of TGFβ signaling proteins in SF3B1 mutant MDS. **(a)** Sashimi plots of bulk RNA-sequencing data of an aberrant 3’ splice site usage in TGFBR1, MAP3K7 and SMURF2 mRNA in SF3B1 mutant acute myeloid leukemia (AML) (top; n=76 patients), SF3B1 wild-type (WT) AML (middle; n=739 patients), and normal bone marrow (bottom; n=26 patients). Red lines indicate SF3B1 mutant-specific junctions while black lines represent junction spanning reads in wild-type cells. The number of reads is listed, and the frequency of reads is in parentheses. **b**, Crystal structure of the short-isoform of TGFBR1 (grey) overlaid with that of the alpha-fold predicted model of SF3B1 mutant induced long-isoform (green). **c,** RT-PCR analysis of aberrant 3’ splice events in *TGFBR1, MAP3K7,* and *SMURF2* in human isogenic K562 cells with knockin of SF3B1^K700E^ mutation. **d**, RT-PCR analysis of endogenous *TGFBR1*, *MAP3K7,* and *SMURF2* splicing in primary samples in healthy bone marrow control patients (n=5), patients with SF3B1^K700E^ mutant MDS (n=5) and non-splicing factor mutant patients with MDS (n=5). **e**, Sanger sequencing electropherogram of the top and bottom PCR products from gel-purified *TGFBR1* RT-PCRs from MDS SF3B1 mutant patients shown in (**c**). The red box highlights the alternatively spliced sequence in the top band, which includes a 12-nucleotide insertion predominantly observed in SF3B1^K700E^ mutant MDS patients. The bottom band sequence is displayed below, showing the canonical exonic sequence. **f**, Western blot of K562 cells with SF3B1 mutation, TGFBR1 knockout (KO), or TGFBR1 KO with addback of TGFBR1 cDNA encoding the long or short isoform. **g,** Western blot of cytoplasmic, membrane, and soluble nuclear fractions of K562 cells with TGFBR1 KO alone or with overexpression of TGFBR1 cDNA encoding the long or short isoform.

**Extended Data Fig. 14:**
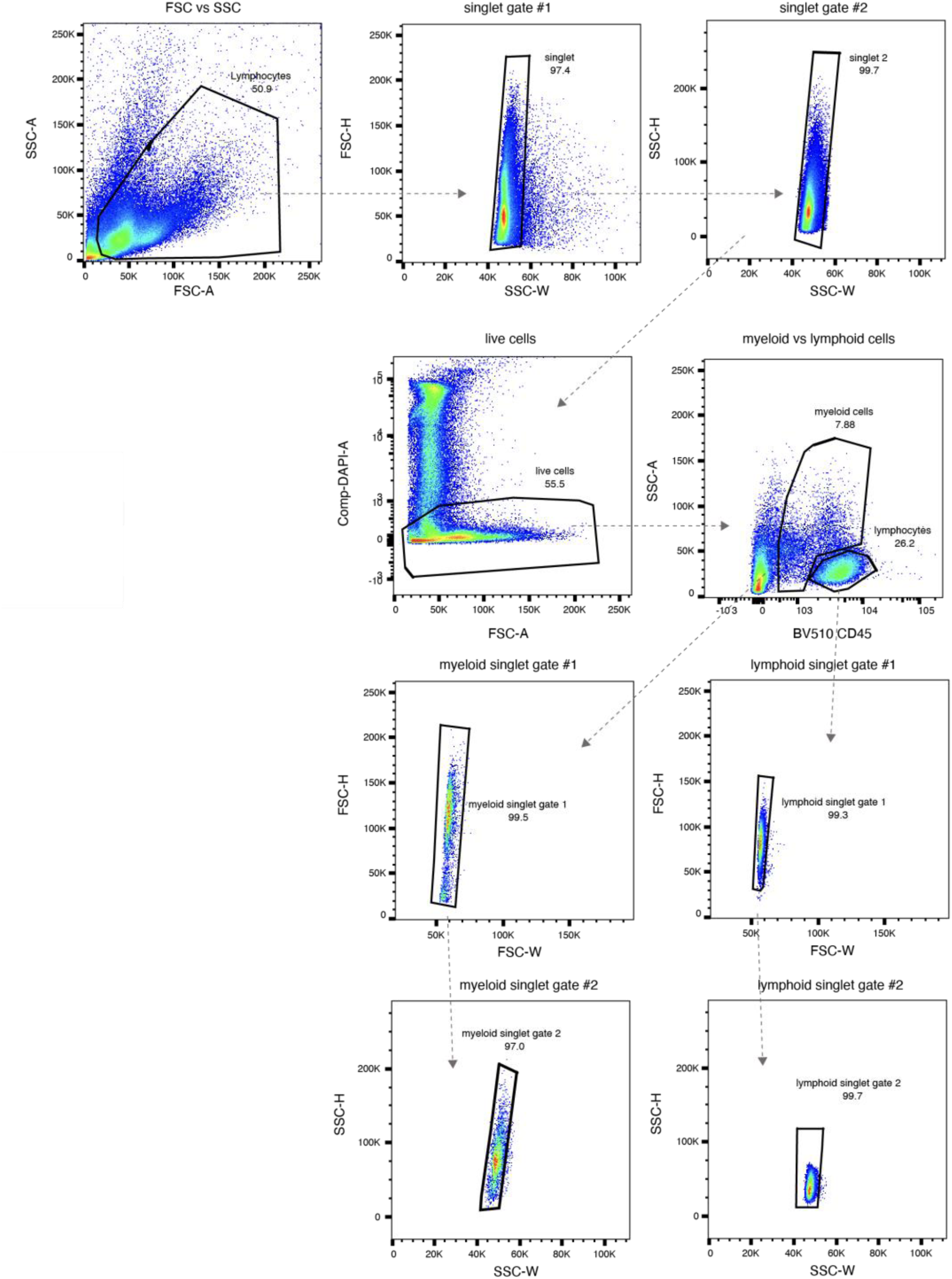
Gating strategy for single cell CITE- and TCR-sequencing studies. Live CD45^+^ lymphocytes and CD45^dim^ non-lymphocytes were sorted for downstream sequencing studies.

